# BOLA3 and NFU1 link mitoribosome iron-sulfur cluster assembly to multiple mitochondrial dysfunctions syndrome

**DOI:** 10.1101/2023.05.27.542581

**Authors:** Hui Zhong, Alexandre Janer, Oleh Khalimonchuk, Hana Antonicka, Eric A Shoubridge, Antoni Barrientos

## Abstract

The human mitochondrial ribosome contains three [2Fe-2S] clusters whose assembly pathway, role, and implications for mitochondrial and metabolic diseases are unknown. Here, structure-function correlation studies show that the clusters play a structural role during mitoribosome assembly. To uncover the assembly pathway, we have examined the effect of silencing the expression of Fe-S cluster biosynthetic and delivery factors on mitoribosome stability. We find that the mitoribosome receives its [2Fe-2S] clusters from the GLRX5-BOLA3 node. Additionally, the assembly of the small subunit depends on the mitoribosome biogenesis factor METTL17, recently reported containing a [4Fe-4S] cluster, which we propose is inserted via the ISCA1-NFU1 node. Consistently, fibroblasts from subjects suffering from “multiple mitochondrial dysfunction” syndrome due to mutations in BOLA3 or NFU1 display previously unrecognized attenuation of mitochondrial protein synthesis that contributes to their cellular and pathophysiological phenotypes. Finally, we report that, in addition to their structural role, one of the mitoribosomal [2Fe-2S] clusters and the [4Fe-4S] cluster in mitoribosome assembly factor METTL17 sense changes in the redox environment, thus providing a way to regulate organellar protein synthesis accordingly.

## INTRODUCTION

The human mitoribosome synthesizes 13 proteins encoded in the mitochondrial genome (mtDNA), which are essential for the transduction of the energy in nutrients into the chemical form of adenosine triphosphate (ATP) by oxidative phosphorylation (OXPHOS) (1,2). Aerobic energy production is critical for life, and alterations in this process due to genetic or environmental perturbations affecting mitoribosome function often result in mitochondrial cardio-encephalomyopathies, neurodegeneration, and certain cancers (2,3). A recent 2.21 Å overall resolution map of the human mitoribosome has uncovered the presence of three [2Fe-2S] clusters, each coordinated by a pair of mitoribosome proteins that provide three and one cysteine thiolates (4). It remains to be determined how these clusters are assembled, whether they play any structural or regulatory role, and how their loss impacts the physiopathology of metabolic disorders resulting from defective Fe-S cluster biogenesis.

Of the three mitoribosomal [2Fe-2S] clusters, one is found in the large subunit (mtLSU) – located in the L7/L12 stalk base, where it bridges the mitochondrion-specific mL66 and the N-terminal extension of uL10m, which provides three and one coordinating cysteine thiolates, respectively (4). In the small subunit (mtSSU), a [2Fe-2S] cluster links mitochondrion-specific extensions of bS18m with three coordinating cysteines and bS6m with one. In the mtSSU lower body, another [2Fe-2S] cluster is coordinated by three cysteine thiolates from the mitochondrion-specific protein mS25 and one from bS16m (4). The three clusters are relatively surface-exposed and located to peripheral proteins (5). Despite their location, the [2Fe-2S] cluster-bridging proteins are essential for mitoribosome function, and some have been linked to human mitochondrial disorders. Specifically, mutations in mS25 lead to encephalomyopathy (6), and mutations in its 2Fe-2S-coordinating partner bS16m caused a syndrome with corpus callosum agenesia, hypotonia, and fatal neonatal lactic acidosis (7).

Another link between Fe-S cluster biogenesis and mitochondrial translation has been recently established. CRISPR-induced loss of the Fe-S cluster biosynthesis protein frataxin (FXN) in human K562 cells resulted in widespread depletion of Fe-S cluster containing proteins as expected but also attenuation of mitochondrial protein synthesis (8). The translation defect was attributed to the loss of the methyltransferase-like protein METTL17, which binds to the mtSSU during late assembly and harbors a previously unrecognized [4Fe-4S] cluster required for the binding of the protein to the mitoribosome (8).

In mammalian cells, the biogenesis and distribution of Fe-S clusters involve two different machineries located in the mitochondria (the iron-sulfur cluster assembly machinery or ISC) and the cytosol (the cytosolic iron-sulfur cluster assembly or CIA) (9–12). *De novo* Fe-S cluster biosynthesis commences in mitochondria with the synthesis of a [2Fe-2S] cluster on a multimeric protein complex, formed by the scaffold protein ISCU2, the cysteine desulfurase complex NFS1/LYRM4/ACP1 that converts Cys into Ala, and FXN (9,13–15). Electrons are also required for the formation of the [2Fe-2S] cluster and are provided by the mitochondrial ferredoxin (FDX2)/ferredoxin reductase (FDXR) system (16,17). The newly synthesized cluster is then released and transferred – with the help of adaptors such as the HSPA9 and HSC20 chaperones (18) – to either a dimer of the monothiol glutaredoxin GLRX5, which inserts it into target [2Fe-2S]-binding proteins or to an apo GLRX5-BOLA3 heterodimeric complex (19,20) for [2Fe-2S] cluster transfer to NFU1 and probably other targets that remain to be identified (21). The [4Fe-4S] clusters are formed by a specific system that includes the ISCA1 and ISCA2 proteins and couples two [2Fe-2S] clusters to generate a [4Fe-4S] cluster (22,23). The [4Fe-4S] clusters are then transferred to target apoproteins directly, as in the case of aconitase (ACO2), or through targeting factors such as NFU1 (9,22,23). NFU1 delivers a [4Fe-4S] cluster to selected apo target proteins with the assistance of additional factors such as NUBPL for ETC complex I (24) or BOLA3 for lipoyl synthase (25).

Mitochondrial proteins harboring Fe-S clusters are involved in essential cellular pathways such as OXPHOS, lipoic acid synthesis, and iron metabolism. Unsurprisingly, there is a long list of disorders associated with Fe-S cluster assembly-machinery factors, such as FXN (Friedreich’s ataxia, (26), ISCU (myopathy with severe exercise intolerance and myoglobinuria (27,28), and GLRX5 (either sideroblastic anemia (29) or nonketotic hyperglycinemia (30)). Also, mutations in proteins involved in the final steps of the maturation of mitochondrial [4Fe-4S]-containing proteins: NFU1 (31,32), BOLA3 (31), IBA57, ISCA2, and ISCA1 were linked to five multiple mitochondrial dysfunction syndromes (MMDS types 1 to 5) (33). Nearly all MMDS subjects present with severe and early onset leukoencephalopathy (33), although certain gene-specific clinical phenotypes such as pulmonary hypertension or dilated cardiomyopathy have been described in MMDS type 1 (*NFU1* mutations) or 2 (*BOLA3* mutations), respectively. In all cases, the mitochondrial phenotypes included combined deficits in ETC enzymes, but defects in mitochondrial translation have not been reported.

To address the biogenesis and physiological significance of the [2Fe-2S] clusters in the mitoribosome, we combined systematic silencing of Fe-S biosynthesis and delivery factors with biochemical and physiological studies in human cultured cells and in fibroblasts from subjects carrying mutations in the Fe-S biogenesis factors BOLA3 and NFU1. Furthermore, we focused on one of the Fe-S coordinating protein pairs to perform structure-guided mutagenesis and dissect the structural and redox-sensing roles of the clusters. Our data unravel the molecular wiring connecting the essential and ancestral mitochondrial function, Fe-S biogenesis, with the control of protein translation in the organelle in health and metabolic disease.

## MATERIALS AND METHODS

### Human cell lines and cell culture conditions

Human HEK293T embryonic kidney cells (CRL-3216, RRID: CVCL-0063) were obtained from ATCC. Control human fibroblasts were obtained from the Montreal Children Hospital Cell bank. Subject fibroblasts and the corresponding over-expression cell lines were described previously (31), and the use of these lines was carried out with approval from The Neuro ethics board.

Cells were cultured in high-glucose Dulbecco’s modified Eagle’s medium (DMEM, Life Technologies) supplemented with 10% fetal bovine serum (FBS), 2 mM L-glutamine, 1 mM sodium pyruvate, 50 μg/ml uridine, and 1x GlutaMAX (Thermo Fisher Scientific) at 37 °C under 5% CO_2_. Cell lines were routinely tested for mycoplasma contamination.

### Key reagents

Tables presenting the list of antibodies, recombinant DNAs, oligonucleotides, and siRNA oligoribonucleotides used in this study are included in **Key Resources Table (Table S2)**.

### Generation of KO cell lines

To create stable human *bS16m*-KO HEK293T cell lines, two gRNA vectors and one linear donor were obtained from OriGene Technologies (KN405837). The gRNAs were designed to target the exon 2 of the *bS16m* locus. The linear donor DNA contained the anti-puromycin gene and was the template for homologous repair (**Supplementary Fig. S1**). The day before transfection, 2.5×10^5^ cells were seeded in a well of the six-well plate, cultured in antibiotic-free media overnight, and replaced it the next day with fresh media. For transfection, 1 μg of one of the gRNA vectors and 1 μg of donor DNA were mixed with 125 μL of Opti-MEM reduced serum media (Gibco, 31985070). Separately, 5 μL of lipofectamine 3000 (Invitrogen, L3000015) were mixed with 125 μL of Opti-MEM reduced serum media and incubated for 5 min at room temperature. Next, the nucleic acid mix and the diluted transfection reagent were combined and incubated at room temperature for 20 min to allow DNA-transfection reagent complexes to form. The complexes were added drop by drop to each well and evenly distributed by rocking the plate gently. The cells were cultured for 2 days in standard conditions, split and diluted 1:10, cultured for 3 days, and then split again 1:10 for up to 7 cycles. Subsequently, the cells were grown in media containing 2.5 μg/mL puromycin (Sigma, P7255) for 7 days, and then diluted to isolate single cells in 96-well plates. Single-cell clones were screened by immunoblotting using a bS16m antibody (Sigma, HPA050081) and genotyping. For genotyping, *bS16m*-forward and *bS16m*-reverse oligonucleotides were used to sequence the *bS16m* locus.

To create stable human *mS25*-KO HEK293T cell lines, 1×10^6^ engineered cell pools were obtained from Synthego (435091–1). Cells were incubated for 2 days, and then isolated as single cells in a 96-well plate. Single cell clones were screened by immunoblotting with an mS25 antibody (Sigma, HPA043490), and then genotyped using oligonucleotides *mS25*-forward and *mS25*-reverse to sequence the *mS25* locus.

The KO cell lines were reconstituted with modified versions of pCMV6-A-Hygro plasmids (OriGene Technologies, PS100024) harboring either wild-type or mutant variants of the corresponding gene under the control of an attenuated CMV6-Δ5 promoter (34). The wild-type human genes *mS25* and *bS16m* were obtained in plasmids from OriGene. Mutations of cysteine/s in bS16m and mS25 coordinating Fe-S clusters to alanine, serine or aspartic acid were generated by engineering the corresponding constructs using Q5 site-directed mutagenesis (New England Biolabs, E0554). For this purpose, 10 pg of template pCMV6-A-Hygro-*bS16m* or Δ5-pCMV6-A-Hygro-*mS25* vector were used, along with the oligonucleotides (**Table S2**), designed to mutate the corresponding codons. After PCR amplification and kinase, ligase & DpnI (KLD) treatment, 2.5 μL of the mixture was transformed into chemically competent *E. coli.* cells (NEB, C2987). Purified plasmids were confirmed by Sanger sequencing.

To transfect the WT or mutant plasmids into KO cell lines, 0.3×10^6^ cells were seeded in one 6-well plate. The next day, 1 μg of plasmids and 5 μL of EndoFectin (GeneCopoeia, EF014) were mixed with 125 μL of Opti-MEM reduced serum media (Gibco, 31985-070) respectively, and then combined. After 20 min incubation, the complexes were added to each well, and cells were cultured in the incubator for 2 days. Subsequently, the cells were split and cultured with media containing 200 μg/mL hygromycin (Invitrogen, 10687010) for 7 days.

### Gene silencing in HEK293T cells

Silencer Select siRNAs for *BOLA3, GLRX5, ISCA1, ISCU, NFS1, NFU1, NUBPL*, and scrambled non-targeting control (NTC) were obtained from Ambion.

Human HEK293T WT cells were seeded in 6-well plates, each well containing 0.15×10^6^ cells. The next day, siRNAs were transfected using lipofectamine RNAiMax (Invitrogen, 13778-150) following the manufacturer’s instructions. Cells were cultured for 3 days, then split and transfected again with the same siRNAs. The process was repeated to reach a total of a 9-day silencing period. The silencing efficiency was confirmed by immunoblotting. The optimal concentrations of siRNAs were determined by titration experiments using different concentrations and silencing for 3 days: 3, 6, 10, 20, 40 nM for *GLRX5, BOLA3, ISCA1*, and *NUBPL*; 5, 20, 50, 100 nM for *NFS1*. The optimal concentrations of siRNAs were used in the 9-day silencing experiments: 40 nM NTC, 10 nM of si*GLRX5*, and a combination of two si*ISCA1* at 20 nM, two si*NFU*1 at 20 nM, two si*BOLA3* at 10 nM, and two si*NUBPL* at 10 nM.

### Whole-cell extracts and mitochondrial isolation

For SDS-PAGE, pelleted HEK293T, KO, siRNA-mediated knockdown and reconstituted cells were solubilized in RIPA buffer (50 mM Tris-HCl pH 8.0, 150 mM NaCl, 1% NP-40, 0.5% sodium deoxycholate, 2 mM EDTA, and 0.1% SDS) with 1 mM phenylmethylsulfonyl fluoride (PMSF) and 1 X EDTA-free mammalian protease inhibitor cocktail (Roche Diagnostics, 11836170001). Cell pellets from control and subject fibroblasts and corresponding overexpression cells were solubilized in 1% n-dodecyl-β-D-maltoside (DDM, Sigma) in PBS with 1x cOmplete™ protease inhibitor cocktail (Roche Diagnostics, 11836170001). Whole-cell extracts were cleared by 15 min centrifugation at 20,000 x *g* at 4 °C.

Mitochondria-enriched fractions from HEK293T cells were isolated from at least ten 80%-confluent 15-cm plates, or 1 L of liquid culture, as described previously (35). Briefly, the cells were collected and resuspended in T-K-Mg buffer (10 mM Tris–HCl, pH 7.4, 10 mM KCl, 0.5 mM MgCl_2_, 4°C) and disrupted with 15 strokes in a pre-chilled homogenizer (Kimble/Kontes). A sucrose stock solution (1 M sucrose, 10 mM Tris–HCl, pH 7.4) was added to the homogenate to reach a final concentration of 0.25 mol/L sucrose. A postnuclear supernatant was obtained by centrifuging the samples twice for 3 min at 1,200 x *g* at 4°C. Mitochondria were pelleted by centrifugation for 10 min at 10,000 x *g* and resuspended in 0.32 M sucrose, 20 mM Tris-HCl, pH 7.4, 1 mM PMSF, and 1x EDTA-free protease inhibitor cocktail.

Mitochondria were also isolated from three near-confluent 15-cm plates of control and subject fibroblasts and corresponding overexpression cells. Pelleted cells were resuspended in ice-cold 250 mM sucrose/10 mM Tris-HCl (pH 7.4) and homogenized with 7 passes as above. A post-nuclear supernatant was obtained by centrifuging the samples twice for 10 min at 600 x *g*. Mitochondria were pelleted by centrifugation for 10 min at 10000 x *g* and washed twice in the same buffer.

### Sucrose gradient sedimentation analysis

Sucrose gradient sedimentation analyses of HEK293T, KO and reconstructed cells were performed as described previously (36). Two mg of mitochondria were extracted in 400 μL of extraction buffer (20 mM Hepes pH 7.4, 100 mM KCl, 20 mM MgCl_2_, 0.50% digitonin, 0.5 mM PMSF, 1x Protease Inhibitor, 40 U RNaseOUT). The lysate was ultracentrifuged at 24,000 x *g* for 15 min at 4 ^°^C. The supernatant was collected and loaded on top of a 5 mL 0.3 M to 1 M sucrose gradient solution (20 mM Hepes pH 7.4, 100 mM KCl, 20 mM MgCl_2_, 0.10% digitonin, 0.5 mM PMSF, 1x Protease Inhibitor, 0.3 or 1 M sucrose, 0.5 mM ribonucleoside vanadyl complex). The gradients were centrifuged at 152,000 x *g* for 3 hours and 10 min at 4 ^°^C, and then fractionated into 15 individual tubes, followed by immunoblotting analysis.

Mitochondria from fibroblasts (200 µg) were lysed in lysis buffer (260 mM sucrose, 100 mM KCl, 20 mM MgCl_2_, 10 mM Tris-Cl pH 7.5, 1% Triton X-100, 5 mM β-mercaptoethanol, cOmplete™ protease inhibitor cocktail without EDTA) on ice for 20 min, centrifuged at 10,000 x *g* for 45 min at 4°C and subsequently loaded on a 1ml 10 - 30% discontinuous sucrose gradient (50 mM Tris-Cl, 100 mM KCl, 10 mM MgCl_2_) and centrifuged at 32,000 rpm for 130 min at 4°C in a Beckman SW60-Ti rotor. After centrifugation, 14 fractions were collected from the top and used for SDS-PAGE analysis.

### *In cellula* pulse-labeling of mitochondrial translation products

*In cellula* mitochondrial protein synthesis was assayed by pulse labeling HEK293T cell lines under study and in control and subject fibroblasts. The cells were cultured in 6-well plates and reached 70-80%-confluent before the experiment. The cells were first incubated in DMEM without methionine for 20 min and then supplemented with 100 µg/mL emetine to inhibit cytoplasmic protein synthesis as described (35). 100 μCi/mL [^35^S]-methionine (PerkinElmer Life Sciences, NEG709A001MC) were added to the cells to label newly synthesized proteins. After 30 min (HEK293T cells) or 60 min (fibroblasts) incubation at 37 °C, the cells were collected by trypsinization, and whole-cell extracts were prepared as mentioned above. Fifty μg of each cellular sample was loaded on a 17.5% (HEK293T cells) or 12-20% gradient (fibroblasts) polyacrylamide gel. Gels were either transferred to a nitrocellulose membrane or exposed to Kodak X-OMAT X-ray film. The membranes were subsequently probed with a primary antibody against β-ACTIN as a loading control or stained with Bio-Safe Coomassie stain (Biorad), dried, and exposed to a Phosphorimager cassette and individual detected bands were quantified using Fiji software (37).

### Cell respiration

Endogenous cell respiration was measured polarographically at 37 °C using a Clark-type electrode (Hansatech Instruments). Cell respiration was assayed in cultured cells, as reported (38). Briefly, approximately 2 x 10^6^ cells were collected and washed with PBS. Pelleted cells were resuspended in 0.5 ml of respiration buffer (RB) containing 0.3 M mannitol, 10 mM KCl, 5 mM MgCl_2_, 0.5 mM EDTA, 0.5 mM EGTA, 1 mg/ml BSA, and 10 mM KH_3_PO_4_, pH 7.4, air-equilibrated at 37 °C. The cell suspension was immediately placed into the polarographic chamber to measure oxygen consumption rates. The specificity of the assay was determined by the cell respiration rates after inhibiting complex IV activity with 0.8 µM KCN. Values were normalized by total cell number.

### Mitoribosome immunoprecipitation from mammalian whole cells

Cells (2.5 × 10^6^) were collected and permeabilized with 2.8 mg/mL digitonin on ice for 10 min. After washing with PBS, pelleted cells were resuspended in 150 μL of lysis buffer (20 mM Hepes pH 7.4, 100 mM KCl, 20 mM MgCl_2_, 0.50% digitonin, 0.5 mM PMSF, 1x Protease Inhibitor, 40 U RNaseOUT) and incubate on ice for 10 min. The extraction was stopped by adding 300 μL of non-detergent buffer. The lysates were centrifuged at 15,000 × *g* for 5 min at 4 °C to obtain clear supernatants. The supernatants were loaded onto washed protein A agarose beads (Thermo Fisher Scientific, 20365) together with 4 μL of IgG (EMD Millipore, 12-370), or a combination of 4 μL each, anti-mS25 (Proteintech, 15277-1-AP) and anti-bL12m (Proteintech, 14795-1-AP) antibodies. The samples were rotated at 4 ^°^C for 4 hours or overnight. The pulled-down proteins were eluted with elution buffer (70 μL of 1% SDS, 62.5 mM Tris-HCl, pH 7.2) and boiled at 96 ^°^C for 5 min.

### Immunoblotting

Protein concentration was measured by the Lowry method (39) or by Bradford assay (40). Protein lysates were resolved by denaturing SDS–PAGE in the Laemmli buffer system (41) and then transferred to nitrocellulose membranes. Membranes were blocked with 5% non-fat milk in western rinse buffer (150 mM NaCl, 10 mM Tris-HCl, pH 8.0, 1 mM EDTA, 0.1% Triton X-100), incubated with primary antibodies overnight at 4 °C, and then secondary antibodies for 1 hour at room temperature. Signals were detected by chemiluminescence incubation and exposure to X-ray film. Optical densities of the immunoreactive bands were measured using the ImageJ software version 1.53r in digitalized images.

### Thiol trapping analyses of mS25, bS16m, uL10m, mL66, bS18m, and METTL17

Reverse thiol trapping assay was used to analyze the *in organello* redox state of cysteines in mS25, bS16m, uL10m, and mL66, as described (42) (43). Briefly, 30 µg of mitochondria purified from WT cell lines were first incubated in translation buffer (60 µg/mL amino acids, 100 mM mannitol, 10 mM sodium succinate, 80 mM KCl, 5 mM MgCl_2_, 1 mM K_3_PO_4_ pH 7.4, 25 mM HEPES pH 7.4, 5 mM ATP, 20 mM GTP, 6 mM phosphocreatine disodium, 60 µg/mL creatine phosphokinase) with either 1 mM H_2_O_2_ or no extra addition for 3 hours at 37 ^°^C. After that, the mitochondria were washed with translation buffer to remove H_2_O_2_. The mitochondria were resuspended in 80 mM membrane-permeable iodoacetamide (IAA), which covalently binds free thiols. After 30 min incubation at room temperature, the samples were washed twice with PBS and then resuspended in 30 µL of PBS. To reduce all the native oxidized cysteines, 5 mM of the reducing agent TCEP (tris(2-carboxyethyl)phosphine) and 2% SDS were added to the samples. The mixtures were incubated at 95 °C for 12 minutes and cooled to room temperature. 4.5 mM AMS (4-acetamido-4′-maleimidylstilbene-2,2′-disulfonic acid) was added to bind all the free cysteines. Finally, the samples were supplemented with Laemmli sample buffer and analyzed by 14% SDS-PAGE followed by immunoblotting with relevant antibodies.

### Iron incorporation assay

The ^55^Fe incorporation assay into the mitoribosomes was done as previously described (44). Briefly, following two transfections (with a 24 h interval) with siRNAs to knockdown *GLRX5*, *BOLA3*, *ISCA1, NFU1,* or *NUBPL* expression, cells were grown in the presence of 1 μM ^55^Fe-transferrin for three days. 2.5×10^6^ cells were collected, followed by whole-cell extract preparation and mitoribosome immunoprecipitation using IgG or a mix of bL12m and mS25 antibodies. The pulled-down proteins were eluted with elution buffer (70 μL of 1% SDS, 62.5 mM Tris-HCl, pH 7.2) and boiled at 96 ^°^C for 5 min. The incorporation of ^55^Fe into mitoribosomes was measured by liquid scintillation counting of eluted pull-down fractions.

### Protein expression in *E. coli*

The plasmids mS25-FLAG_pET-21a, bS16m-HA_pET-21a, mS25-Flag_bS16m-HA-His_pETDuet-1 were transformed into four competent *E. coli* strains: Rosetta (Novagen, 71402-3), One Shot BL21(DE3) (Invitrogen, 44-0184), OverExpress C43 (DE3) (Biosearch, 60446-1), and BL21 (DE3) pLysS (Promega, L1195) following the manufacturer’s instructions. For protein expression and solubility trials, the bacterial cultures were grown overnight in a shaker-incubator at 30 ^°^C, 25 ^°^C, or 16 ^°^C. The cells were then harvested, resuspended in 1 mL of lysis buffer (20 mM K_3_PO_4_ pH 8, 1 M KCl, 0.2 mM PMSF, 1 g/L lysozyme), and incubated at 37 ^°^C for 1 h. Following sonication for 3 cycles of 30 seconds with cooling intervals, at intensity 12 in an ultrasonic cell disrupter (Virsonic 100), the lysates were centrifuged at 18,000 *x g* for 30 min at 4 ^°^C. The supernatants were collected, and the pellets were resuspended with 450 mL of ddH_2_O. Both the supernatants and pellets were analyzed by immunoblotting. For immunoprecipitation experiments, mS25-Flag_bS16m-HA-His_pETDuet-1 plasmids were transformed into OverExpress C43 (DE3) competent cells and cultured overnight at 30 ^°^C. Cells were harvested and lysed as described above. Anti-FLAG-conjugated agarose beads were added to the supernatants and incubated in an orbital rotator at 4 ^°^C overnight. The proteins bound to the beads were eluted with 2x Laemmli buffer.

### Blue-Native PAGE (BN-PAGE)

BN-PAGE was used to separate individual OXPHOS complexes (45). Fibroblasts were pelleted from near-confluent 10-cm plate and resuspended in PBS to achieve a final protein concentration of 5 mg/ml. Digitonin was added to the cells at a ratio of 0.8 mg digitonin/mg of protein, and the samples were incubated on ice for 5 min, diluted to 1.5 ml with PBS, and centrifuged for 10 min at 10,000 x *g*, at 4 ^°^C. The pellet was resuspended in a buffer containing 75 mM Bis-Tris (pH 7.0), 1.75 M aminocaproic acid, and 2 mM EDTA at a final protein concentration of 2 mg/mL. 1/10 volume of 10% n-dodecyl β-D-maltoside was added to the samples, and the samples were incubated on ice for 30 min, followed by centrifugation at 20,000 x *g* for 20 min, at 4 ^°^C. The supernatant (20 µg) was run on 6–15% polyacrylamide gradient gels as described in detail elsewhere (46). Separated proteins were transferred to a nitrocellulose membrane using a semi-dry system (BioRad), and immunoblot analysis was performed with the indicated antibodies. For loading control, BN-PAGE samples were supplemented with SDS-loading buffer, denatured, run on an SDS-PAGE, and immunoblotted with indicated antibodies.

### Isolation of genomic DNA from mammalian cells

Genomic DNA was extracted from HEK293T WT, *bS16m*-KO, and *mS25*-KO cell lines as previously described (47) with slight modifications. Briefly, the cell pellets were suspended in RSB buffer (10 mM Tris-HCl, pH 7.5, 10 mM NaCl, 25 mM EDTA pH 8, 1% SDS, 1 mg/mL proteinase K), and then incubated at 55 °C for 2 hours. 0.1 mg/mL of RNAse was added and incubated at 37 °C for 15 min to remove RNA. The DNA was further extracted with phenol/chloroform /isoamyl alcohol buffer and precipitated with ethanol and sodium acetate.

### Mitochondrial DNA levels determination by quantitative RT-PCR

To estimate mtDNA levels in HEK293T WT, *bS16m*-KO, and *mS25*-KO cell lines, DNA samples were analyzed by quantitative Real Time (RT)-PCR using SYBR Green supermix (Bio-Rad, 1725271), following standard procedures in a CFX96/C1000 Real-Time PCR Detection System (Bio-Rad). Quantitative Real-Time PCR was performed according to Minimum Information for Publication of Quantitative Real-Time PCR Experiments (MIQE) guidelines (48). A standard curve was generated for each pair of primers, and efficiency was measured between 90%-110%. We used the comparative cycle threshold (Ct) method to determine the relative quantity of mtDNA. The calculations of mtDNA levels in each sample were performed using the cycle threshold (CT) values (also known as quantification cycle (Cq), according to the RDML (Real-Time PCR Data Markup Language) data standard (49), and the ΔΔCt method (50), using *ND2* levels as a mtDNA indicator, and *ACTIN* DNA levels as the internal control. Three independently cultured biological replicates (performed on different days with freshly made reagents) each with three technical replicates (three measurements per sample), were analyzed. Comparisons among groups were performed using one-way ANOVA with post-hoc Tukey HSD test. Each group was compared with the WT sample. Statistical significance was established at p < 0.05. The sequences of primers used in this study (*ND2*_F and *ND2*_R, *ACTIN*_F and *ACTIN*_R) are listed in **Table S2**.

### TMT quantitative proteomics and mass spectrometry

Tandem mass tags (TMT), coupled with liquid chromatography–tandem mass spectrometry (LC–MS/MS) proteomic analysis, were used to investigate the proteomic profiles of HEK293T cells treated for 9 days with siRNA oligonucleotides to silence the expression of *GLRX5, BOLA3, NFU1, NBUPL*, or a non-targeting (NT) control. Following silencing, whole cell pellets were collected and sent to the Proteomics Core at the University of Florida-Scripps Biomedical Research.

Samples in 5% SDS (v/v) were brought to 24 μL with an additional 5% SDS and processed for digestion using micro-S-Traps (Protifi) according to the manufacturer’s instructions. Briefly, proteins in 5% SDS were reduced with 1 μL of 120 mm TCEP [Tris (2-carboxyethyl) phosphine hydrochloride] at 56 ^°^C for 20 min, followed by alkylation using 1 μL of 500 mM methyl methanethiosulfonate for 20 min at ambient temperature. Finally, 4 μg of sequencing-grade trypsin was added to the mixture and incubated for 1 h at 47 °C. Following this incubation, 40 μL of 50 mm TEAB (tetraethylammonium bromide) was added to the S-Trap, and the peptides were eluted using centrifugation. Elution was repeated once. A third elution using 35 μL of 50% acetonitrile was also performed, and the eluted peptides were dried under a vacuum. The peptides were subsequently resolubilized in 30 μL of 50 mM triethylammonium bicarbonate, pH 8.5, labeled with TMT labels (10-plex) according to the manufacturer’s instructions (Thermo Fisher Scientific), and pooled. The multiplexing strategy is provided in **Table S1.** The pooled, plexed samples were then dried under vacuum, resolubilized in 1% trifluoroacetic acid, and finally desalted using 2 μg capacity ZipTips (Millipore) according to manufacturer instructions. Peptides were then eluted into a Fusion Tribrid Mass Spectrometer (Thermo Fisher Scientific) from an EASY PepMap RSLC C18 column (2 μm, 100 Å, 75 μm x 50 cm; Thermo Fisher Scientific), using a gradient of 5–25% solvent B (80:20 acetonitrile/water; 0.1% formic acid) for 180 min, followed by 25–44% solvent B in 60 min, 44– 80% solvent B in 0.1 min, a 5 min hold of 80% solvent B, a return to 5% solvent B in 0.1 min, and finally a 10 min hold of solvent B. All flow rates were 250 nL/min delivered using a nEasy-LC1000 nano-liquid chromatography system (Thermo Fisher Scientific). Solvent A consisted of water and 0.1% formic acid. Ions were created at 1.7 kV using an EASY Spray source (Thermo Fisher Scientific) held at 50 °C.

### Proteomic data processing and statistical analysis

Quantitative analysis of the TMT-MS experiments was performed simultaneously to protein identification (one-step method) using the Proteome Discoverer 2.5 software (Thermo Fisher Scientific). The precursor and fragment ion mass tolerances were set to 10 ppm, 0.6 Da, respectively, enzyme was Trypsin with a maximum of 2 missed cleavages and UniprotKB Human and common contaminant FASTA file used in SEQUEST searches. The impurity correction factors obtained from Thermo Fisher Scientific for each kit were included in the search and quantification. The following settings were used to search the data: dynamic modifications; Oxidation / +15.995Da (M), Deamidated / +0.984 Da (N, Q), and static modifications of TMTpro / +304.207 Da (K) (N-Terminus, K), MMTS / +45.988 Da (C). Scaffold Q+ (version Scaffold_5.0.0, Proteome Software Inc.) was used for TMTpro peptide and protein identifications. Peptide identifications with >82.0% probability were accepted to achieve an FDR less than 1.0% by the Percolator posterior error probability calculation (51). Proteins with >95.0% probability were accepted to achieve an FDR less than 1.0% and contained at least 1 identified peptide. Protein probabilities were assigned by the Protein Prophet algorithm (52), and default Scaffold settings were used except for Blocking level was Unique Samples, means were used for averaging, and normalization was applied as reported (53). Of 40824 spectra in the experiment at the given thresholds, 33826 (83%) were included in the quantitation. Differentially expressed proteins were determined by applying ANOVA with an unadjusted significance level of p < 0.05, corrected by Benjamini-Hochberg.

### General statistical analysis

Unless indicated otherwise in the figure legends, all experiments were performed at least in biological triplicates, and results are presented as mean ± standard deviation (SD) or standard error of the mean (SEM) of absolute values or percentages of control. Statistical *p* values for the comparison of two groups were obtained by applying a Student’s two-tailed unpaired *t*-test. For the comparison of multiple groups, one-way analysis of variance (ANOVA) followed by Tukey’s post-hoc test or two-way ANOVA with a Dunnett correction for multiple comparisons were performed. A *p* < 0.05 was considered significant. The *p* values are indicated in the graphs: * *p* < 0.05, ** *p* < 0.01, *** *p* < 0.001, and **** *p* < 0.0001.

## RESULTS

### The [2Fe-2S] cluster-coordinating cysteines in mitoribosomal proteins mS25 and bS16m are crucial to their stability and structural function

To study the role of [2Fe-2S] clusters and their coordinating proteins in the assembly and function of the mitoribosome, we focused on the Fe-S cluster-binding pair bS16m-mS25 (**Fig. 1A**), two proteins involved in mitochondrial disorders (6,7). In the mtSSU lower body, the [2Fe-2S] cluster is coordinated by three cysteine thiolates from the mS25 (C139, C141, and C149) and one from bS16m (C26) (**Fig. 1A**, (4)).

**Figure 1.**
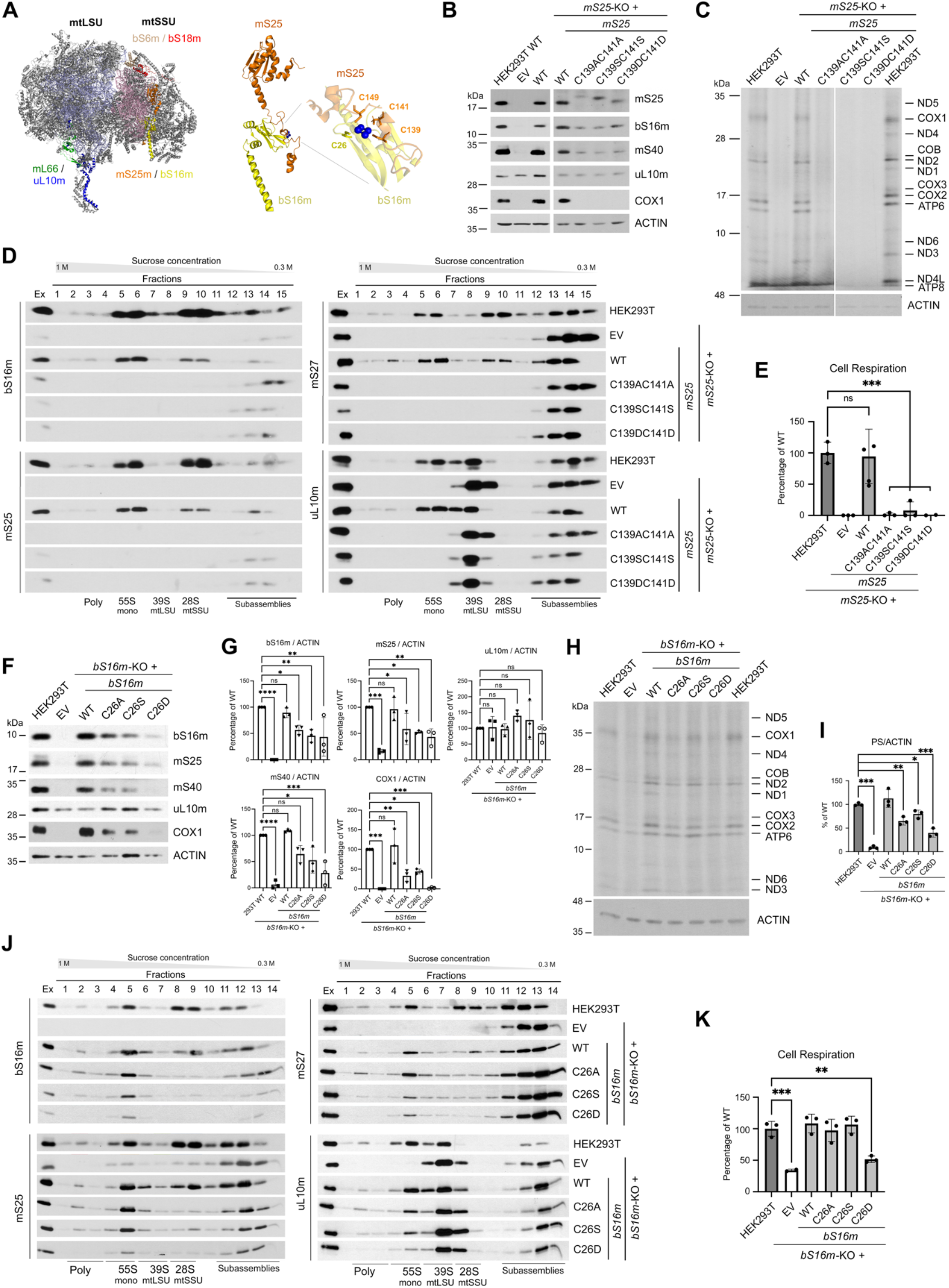
The [2Fe-2S] cluster-coordinating cysteine residues in mS25 and bS16m are essential for mitoribosome assembly and function. (**A**) Mitoribosome cryo-EM reconstitution (PDB: 7qi4) in which the pairs of proteins that coordinate a [2Fe-2S] cluster are indicated. The right-side panel shows the mS25-bS16m pair magnified, and the residues involved in [2Fe-2S] cluster coordination are indicated. mS25 residues are shown in orange, and bS16m are shown in yellow. The [2Fe-2S] cluster is depicted as 4 blue spheres. Figures were prepared in Pymol and Adobe Photoshop. (**B, F**) Steady-state levels of mS25, bS16m, additional mitoribosome markers (mS40 and uL10m), and a marker of mitochondrial protein synthesis (complex IV subunit COX1) were assessed by immunoblotting in HEK293T wild-type cells, and *mS25*-KO (**B**) or *bS16m*-KO cells (**F**) reconstituted with an empty vector (EV), or plasmids expressing the corresponding wild-type protein (WT), or variants carrying mutations in the indicated [2Fe-2S] cluster coordinating cysteines. ACTIN was used as a loading control. (**C, H**) Mitochondrial protein synthesis following ^35^S-methionine incorporation into newly synthesized mitochondrial proteins in the presence of emetine to inhibit cytoplasmic protein synthesis. ACTIN was used as a loading control. (**D, J**) Sucrose gradient sedimentation analyses of mtSSU (bS16m, mS25, and mS27) and mtLSU markers (uL10m) in mitochondria purified from the indicated cell lines and extracted in the presence of 100 mM KCl, 20 mM MgCl_2_, and 0.5% digitonin. (**E, K**) KCN-sensitive endogenous cell respiration for indicated cell lines measured polarographically. The bar graphs represent the average ± SD of three independent experiments. Black dots represent individual data points. Two-tailed unpaired *t*-test, \*\**p* < 0.01; \*\*\**p* < 0.001; n.s., not significant. (**G, I**) The graphs show the densitometry (average ± SD) of three independent experiments as in panel F (**G**) or H (**I**). Two-tailed unpaired *t*-test, \**p* < 0.05, \*\**p* < 0.01, \*\*\**p* < 0.001, \*\*\*\**p* < 0.0001. In panel I, *PS* indicates protein synthesis.

Using the CRISPR-Cas9 technology, we created stable human HEK293T cell line KOs for *mS25*, or *bS16m,* which maintained WT mtDNA levels (**Supplementary Fig. S1**). The stability of mS25 and bS16m depended on one another, and the assembly of the mtSSU depended on both (**Fig. 1B-K**). However, whereas mS25 is essential for mtSSU biogenesis (**Fig. 1D**), the absence of bS16m allows for a small residual amount of 28S mtSSU, and 55S monosome assembly (**Fig. 1J**) that supports ∼8% of protein synthesis (**Fig. 1H, I**) and 25% of cellular respiratory capacity compared to wild-type (WT) HEK293T cells (**Fig. 1K**). Therefore, mS25 can assemble – albeit inefficiently – in the absence of bS16m, but not *vice versa*. All the phenotypes in the KO lines were restored to WT levels by rescue with the corresponding WT genes (**Fig. 1B-K**), indicating a lack of CRISPR off-target effects. To assess the role of [2Fe-2S]-coordinating residues in mS25, we reconstituted the *mS25*-KO cell line with mS25 variants in which the C139-C141 cysteine pair was replaced with serine, alanine, or aspartic acid residues. Although each of these variants failed to rescue mtSSU assembly and function (**Fig. 1B-E**), the three proteins accumulated, albeit at levels lower than the WT, and nearly equally supported the stability of bS16m (**Fig. 1B**). These data suggest that mS25 and bS16m may associate independently of the Fe-S cluster. To assess the role of the [2Fe-2S]-coordinating residue in bS16m, we reconstituted the *bS16m*-KO cell line with bS16m variants in which C26 was replaced with serine, alanine, or aspartic acid. The three bS16m proteins were also expressed and accumulated at lower than the WT levels and stabilized proportional amounts of mS25 (**Fig. 1F**). Furthermore, these variants supported mtSSU and monosome assembly (**Fig. 1J**) and function (**Fig. 1H-I**) – protein synthesis was rescued up to ∼50% of WT levels for the C26A and C26S variants, and to ∼25% for the C26D mutant, which resulted in full or 50% restoration of respiration, respectively (**Fig. 1K**). These data suggest that either unstable coordination of the [2Fe-2S] cluster by the three sulfhydryl groups in mS25 or the complete lack of the cluster due to the absence of bS16m coordinating cysteine, markedly attenuates mitoribosome function.

The interconnected lability of mS25 and bS16m suggests that their shared [2Fe-2S] cluster facilitates structural stability thereof. Considering this, three potential scenarios could account for their assembly into the mitoribosome. The pair of proteins could incorporate into the mitoribosome sequentially and subsequently, acquire the cluster. Alternatively, they could interact first, incorporate into the mitoribosome, and then acquire the cluster. Finally, these subunits could incorporate into the assembling mitoribosome as pre-formed [2Fe-2S]-bound heterodimers. To examine these scenarios, we expressed mS25 and bS16m in *E. coli* and analyzed their solubility and association. When bS16m-HA or mS25-FLAG were expressed individually from the pET-21a plasmid, either protein was completely insoluble (not shown). However, when the two proteins were co-expressed from the pETDuet-1 plasmid, they remained partially soluble, either when the WT or C139AC141A mS25-FLAG variants were co-expressed with WT bS16m-HA. Immunoadsorption assays using the soluble fractions and anti-FLAG-conjugated beads efficiently co-purified mS25 and bS16m (**Supplementary Fig. S2A-C**). These data indicate that mS25 can associate with bS16m, irrespective of their shared Fe-S cluster coordination.

### The mitoribosome [2Fe-2S] cluster coordinated by mS25 and bS16m acts as a redox sensor to attenuate mitochondrial translation under oxidative stress

Some [Fe-S] cluster proteins are known to sense and regulate gene transcription/translation in response to environmental stimuli (54). The [2Fe-2S]-containing sensors can lose their cluster or modify their redox state to effect conformational changes that modulate their activity (54). The peripheral locations of the mitoribosomal [2Fe-2S] clusters suggest a potential role in oxidative stress sensing. To assess the effect of oxidative stress on mitoribosome function, we analyzed mitoribosome protein synthesis by following the incorporation of ^35^S-methionine into newly synthesized polypeptides in the presence of emetine to inhibit cytosolic protein synthesis. Upon treatment with 0.25mM or 0.50 mM H_2_O_2_ for 3 hours, which induces mild oxidative stress in cells (55), the mitoribosome-driven protein synthesis was decreased down to ∼40% and ∼15% of untreated cells, respectively (**Fig. 2A**), whereas the mitoribosome proteins remained stable (**Supplementary Fig. S3**). These data suggest that loss or remodeling of Fe-S cluster-binding in mitoribosomal proteins could contribute to the effect of oxidative stress on mitoribosome function.

**Figure 2.**
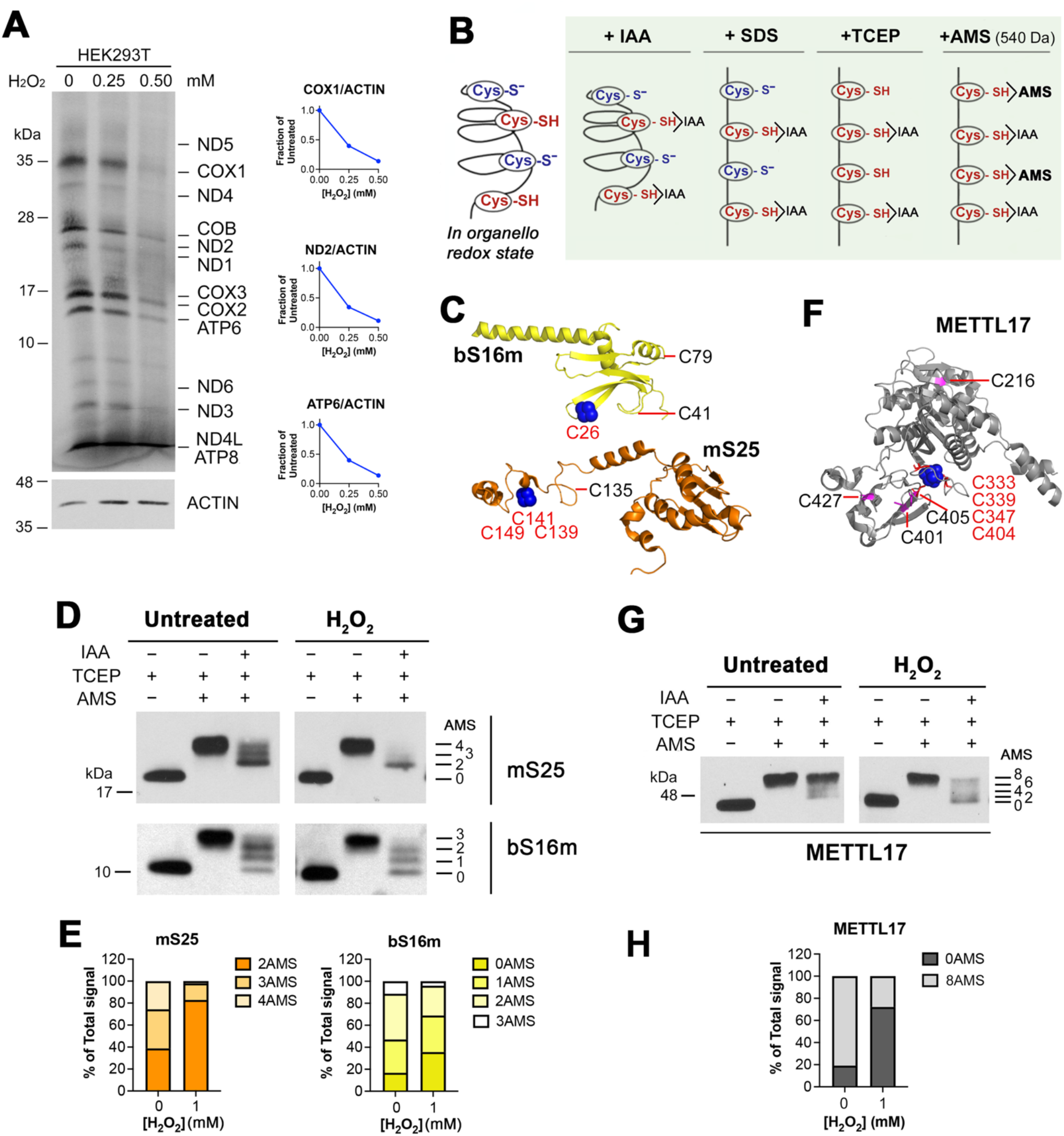
Cysteines in [2Fe-2S] cluster-coordinating proteins are sensitive to oxidative stress. (**A**) Mitochondrial protein synthesis following ^35^S-methionine incorporation into newly synthesized mitochondrial proteins in the presence of emetine to inhibit cytoplasmic protein synthesis. Wild-type HEK293T cells were treated with 0, 0.25, or 0.50 mM H_2_O_2_ for 3 h. ACTIN served as a loading control. The right-side graphs represent the quantification of the signal in a digitalized image using the Adobe Acrobat Histogram tool. The data for COX1, ND2, and ATP6, normalized by ACTIN and expressed as the fraction of untreated cells, are presented. (**B**) Reverse thiol trapping approach to detect the native cysteine residues inaccessible to a cell-permeable alkylating compound (2-iodoacetamide, IAA) that upon denaturation with SDS (sodium dodecyl sulfate) and full reduction with TCEP (tris(2-carboxyethyl)phosphine) are bound to AMS (4-acetamido-4’-maleimidylstilbene-2,2’-disulfonic acid), which adds ∼540 Da per thiol group. (**C** and **F)** Cryo-EM structures of the mS25-bL16m pair of proteins that coordinate a [2Fe-2S] cluster in the mtSSU (PDB: 7qi4) (**C**), and the mtSSU assembly factor METTL17 that harbors a [4Fe-4S] cluster (PDB: 8csp) (**F**) The [Fe-S] cluster-coordinating cysteine residues are indicated in red, and all additional cysteines in each protein are depicted in black. The Fe-S clusters are shown (not to scale) as blue spheres. Figures were prepared in Pymol and Adobe Photoshop. (**D** and **G)** Reverse thiol trapping of mS25 and bS16m (**D**) or METTL17 (G) in mitochondria isolated from WT HEK293T cells, treated or not with 1 mM H_2_O_2_ for 3 h. (**E** and **H)** The bar graphs represent the number of AMS-bound cysteines (unavailable to IAA binding *in organello*). The data are the average of two independent experiments. Figures are representative of two independent repetitions with similar results.

If the mitoribosome [2Fe-2S]-coordinating proteins lost their cluster in response to oxidative stress, the coordinating cysteines could become accessible for modification. To evaluate this scenario, we used iodoacetamide (IAA)/ 4-acetamido-4′-maleimidylstilbene-2,2′-disulfonic acid (AMS)-based reverse thiol trapping approach (**Fig. 2B**) to assess the redox state of the pairs mS25-bS16m and mL66-uL10m in functional mitochondria isolated from HEK293T cells treated or not with H_2_O_2_. Mitochondria were first treated with translation buffer containing an excess of membrane-permeable IAA, which covalently binds to all free thiols. Next, the samples were denatured and reduced using sodium dodecyl sulfate (SDS) and tris-[2-carboxyethyl] phosphine (TCEP), respectively. The cysteines protected from IAA were subsequently identified by testing molecular size after incubating the samples with AMS, which can bind to reduced cysteines, adding 540 Da per molecule to the proteins of interest.

mS25 has four cysteines, three of which are engaged in [2Fe-2S] cluster coordination (**Fig. 2C**). Under standard conditions, mS25 accumulated in three distinct populations: 40% of the total protein exhibited two cysteines inaccessible by IAA, 35% presented with three (3-AMS), and 25% carried four (4-AMS) modified groups (**Fig 2. D-E**). Upon H_2_O_2_ treatment, this ratio shifted to 82% of mS25 with two cysteines inaccessible by IAA, and 18% with three modified thiolates. In this setting, only the fractions with 3- and 4-AMS groups could have three cysteines coordinating the [2Fe-2S] cluster. Their contribution shifts from 60% of the total in standard conditions to 18% under oxidative stress conditions. bS16m has three cysteines, one of which is engaged in [2Fe-2S] cluster coordination (**Fig. 2C**). Under standard conditions, we observed the accumulation of bS16m in four populations: 15% of the total protein had no cysteines inaccessible by IAA, 30% had one (1-AMS), 45% had two (2-AMS), and 10% had three (3-AMS) modified groups (**Fig. 2D-E**). Following H_2_O_2_ treatment, the proportions shifted to 35%, 35%, 25%, and 5%, respectively (**Fig. 2D-E**). All fractions with 1 or more AMS groups could potentially coordinate the [2Fe-2S] cluster. The IAA inaccessibility of cysteines in native proteins could also be due to interactions with rRNA. At any rate, the total amount of [2Fe-2S] cluster-ligating thiolates is decreased in the presence of H_2_O_2_ (**Fig. 2D-E**).

In contrast to the mS25-bS16m pair, the [2Fe-2S] cluster coordinated by mL66-uL10m may not have a major role in redox sensing. Four of the five cysteines present in mL66 – which provides three sulfhydryl groups for [2Fe-2S] cluster coordination – were equally inaccessible to IAA under standard and oxidative stress conditions (**Supplementary Fig. S4A-C**). Its partner subunit, uL10m, which harbors four cysteines and provides the fourth coordinating group, accumulated in two populations: 2- and 3-AMS (**Supplementary Fig. S4A-C**). The IAA inaccessibility ratio for uL10 changed from the standard (40%/60%) to oxidative stress conditions (60%/40%). However, the lack of appreciable H_2_O_2_ effect on mL66 suggests only a limited role for the [2Fe-2S] cluster in mL66-uL10m in translation regulation. Similarly, although we could not test the effect of H_2_O_2_ on the redox state of bS6m due to the lack of a properly working antibody, its partner bS18m (contains five cysteines, three of which are coordinating a [2Fe-2S] cluster) did not exhibit any appreciable changes in redox state upon H_2_O_2_ treatment (**Supplementary Fig. S4D-F**).

We then examined the effect of H_2_O_2_ treatment on the [4Fe-4S]-containing mtSSU assembly factor METTL17. The mature protein has 8 cysteines, four of which are engaged in cluster coordination (**Fig. 2F-H**). The protein remained stable upon treatment (**Supplementary Fig. S3**). However, the thiol trapping assay showed that the percentage of molecules with eight (8-AMS) inaccessible cysteines shifted from ∼80% in standard conditions to less than 20% upon H_2_O_2_ treatment. In the latter condition, ∼80% of METTL17 molecules had all eight cysteines free, indicating loss of their coordinating [4Fe-4S] cluster.

We conclude that under oxidative stress conditions, the insult is sensed directly by the existing mitoribosomes via the [2Fe-2S] cluster coordinated by mS25-bS16m. At the same time, the stress is effectively sensed by METTL17 to inhibit the assembly of new mtSSU particles.

### The [Fe-S] cluster delivery nodes GLRX5-BOLA3 and ISCA1-NFU1 support mitoribosome biogenesis

Following the identification of [2Fe-2S] clusters in the mitoribosome, one fundamental query is how they are transferred to recipient apo-mitoriboproteins. We approached this question by examining the effect of systematic 9-day siRNA-mediated depletion of key [Fe-S] cluster biosynthesis and delivery proteins on mitoribosome assembly and function by proteomics and metabolic labeling approaches (**Supplementary Fig. S5A**). Specifically, we targeted the [Fe-S] biosynthetic proteins NFS1, ISCU, and the [2Fe-2S] assembly and delivery proteins GLRX5 and BOLA3. Because mtSSU formation involves the [4Fe-4S] protein METTL17, we also included the [4Fe-4S] cluster assembly and delivery proteins ISCA1, NFU1, and NUBPL. Following trials to determine the optimal siRNA sequence and concentration, we reached, in all cases, a silencing efficacy of >95% as assessed by immunoblotting (**Supplementary Fig. S5B-G**). Using this candidate approach, we confirmed expected attenuations in known [2Fe-2S] client proteins, such as the Rieske iron-sulfur protein (RISP) of OXPHOS Complex III, and [4Fe-4S]-carrying proteins such as aconitase (ACO2), or proteins that contain both species such as the NADH-ubiquinone oxidoreductase 75 kDa subunit (NDUFS1) of Complex I, or the succinate dehydrogenase subunit B (SDHB) of Complex II (**Supplementary Fig. S5B-G**). Silencing of *NFS1, ISCU, GLRX5, BOLA3, NFU1,* and *ISCA1*, but not *NUBPL,* attenuated the levels of mitoribosome markers as well as METTL17 (**Supplementary Fig. S5B-G**).

To unequivocally identify mitochondrial proteins whose stability is regulated by GLRX5, BOLA3, NFU1, or ISCA1, we carried out quantitative tandem mass tag (TMT)-based proteomics on whole cell lysates from HEK293T cells treated with gene-specific siRNAs as indicated above. The results – summarized in four volcano plots (**Fig. 3A**) and two heatmaps (**Fig. 3B**) – indicate that compared to control non-targeting siRNA (siNT)-treated HEK293T cells, many components of the mitoribosome SSU and LSU were attenuated in response to GLRX5 and especially BOLA3 depletion. By contrast, silencing of ISCA1 and NFU1 only decreased mtSSU components but had no effect on the mtLSU stability. Despite the seemingly minor decrease in mitoribosome components, the biological importance of their attenuation is substantiated by its similarity to the attenuation observed in established [2Fe-2S] and [4Fe-4S] client proteins (**Fig 3A**). Thus, our data are congruent with a scenario in which the GLRX5-BOLA3 pathway directly delivers [2Fe-2S] clusters to the two mitoribosome subunits. Furthermore, downstream of GLRX5-BOLA3, the ISCA1-NFU1 pathway targets the [4Fe-4S] cluster to the mtSSU assembly factor METTL17. Cryo-EM structures of human mtSSU assembly intermediates with METTL17 bound to the cleft between the head and the body contain the protein pairs bS16m-mS25 and bS6m-bS18m already coordinating their [2Fe-2S] cluster (56). Therefore, the BOLA3-mediated delivery of [2Fe-2S] clusters to the mtSSU must occur upstream of METTL17.

**Figure 3.**
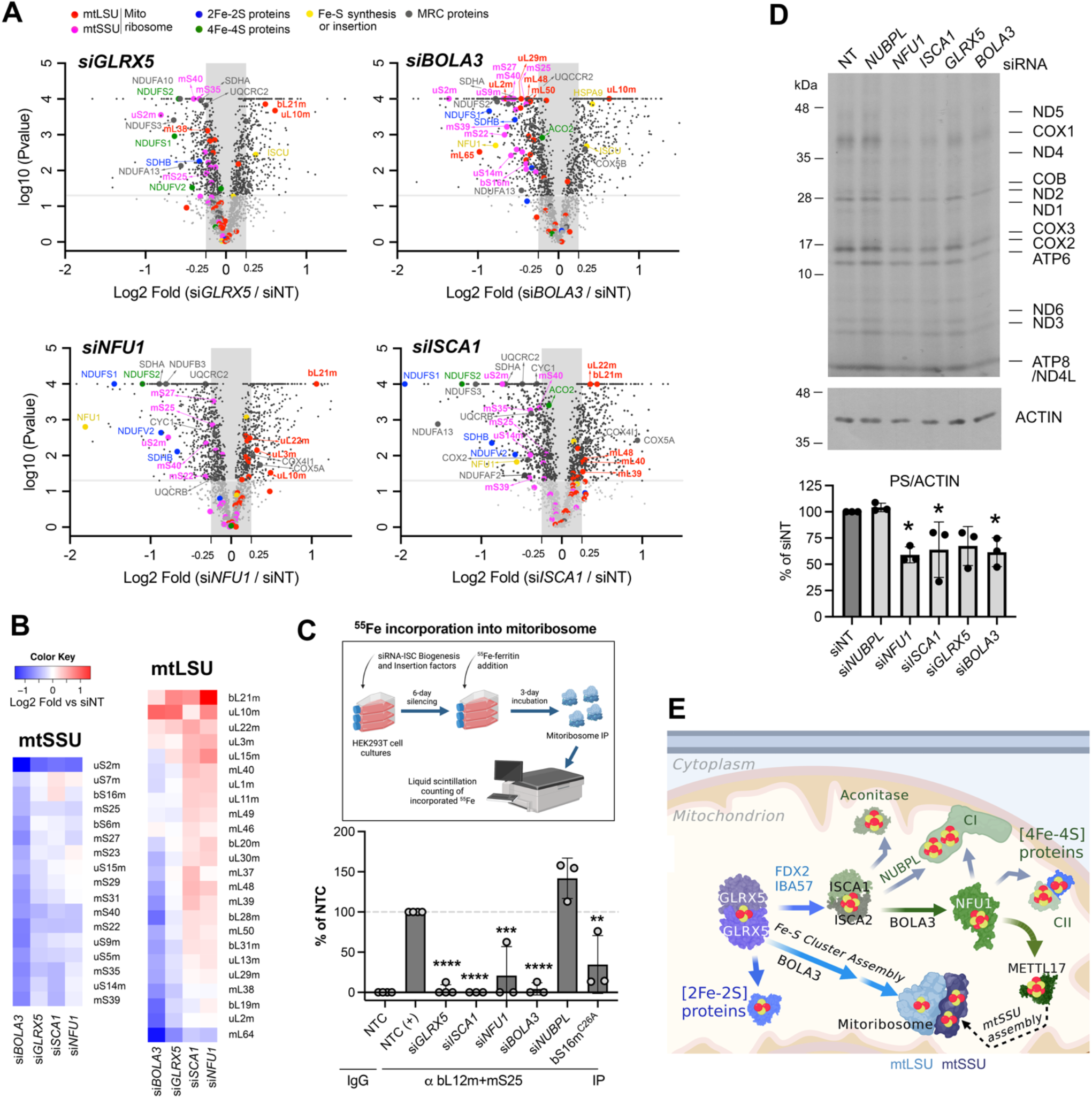
The GLRX5-BOLA3 and ISCA1-NFU1 Fe-S cluster delivery pathways are required for mitoribosome biogenesis. (**A**) Volcano plots displaying differentially expressed proteins between HEK293T cells treated with siNT (non-targeting) and si*GLRX5*, si*BOLA3*, si*ISCA1,* or si*NFU1* for 9 days. The *y*-axis displays the log10 (P values), and the *x*-axis displays the Log2 (fold-change value). The horizontal grey line represents the 10^-1.3^ threshold on the p values, while the vertical lines show thresholds of ± 1.2-fold changes. The raw Multiplex TMT-MS/MS data have been uploaded to PRIDE repository, with ref# 1-20230106-165123. (**B**) Heat map showing the levels of mtSSU and mtLSU proteins in si*GLRX5*, si*BOLA3*, si*ISCA1,* and si*NFU1* cells, normalized by siNT-treated HEK293T cells after 9 days of silencing. The map represents the average of four independent experiments. (**C**) The schematic depicts the approach followed to assess ^55^Fe incorporation into mitoribosomes. The bar graph represents the average ± SD of three independent experiments. Grey dots represent individual data points. Two-tailed unpaired *t*-test, \**p* < 0.05; \*\*\*\**p* < 0.0001 (**D**) Mitochondrial protein synthesis following ^35^S-methionine incorporation into newly synthesized mitochondrial proteins in the presence of emetine to inhibit cytoplasmic protein synthesis. ACTIN was used as a loading control. The bar graph represents the average of two independent experiments. Black dots represent individual data points. (**E**) Schematic showing the delivery of Fe-S clusters in mammalian cells, highlighting the GLRX5-BOLA3 pathway involved in [2Fe-2S] cluster delivery to the mitoribosome, and the ISCA1-NFU1 node that targets a [4Fe-4S] cluster to the mtSSU assembly factor METTL17.

To examine the effects of knocking down [Fe-S] cluster delivery proteins on mitoribosome maturation using a complementary approach, we performed metabolic labeling of HEK293T cells with ^55^Fe and analyzed iron incorporation into mitoribosomes by immunoprecipitation (**Supplementary Fig. S2D-E**) and liquid scintillation counting (**Fig. 3C**). We first tested the ability to immunoprecipitate mitochondrial ribosomes from whole cells extract using antibodies against endogenous proteins mS25 and bL12m (**Supplementary Fig. S5H-I**) and used both together to determine the ^55^Fe incorporation into the mitochondrial ribosome. Upon knockdown of *GLRX5, BOLA3, or ISCA1* in WT cells for five days, the ^55^Fe signal of the mitoribosome was virtually undetectable (**Fig. 3C**). In contrast, NUBPL silencing did not produce any appreciable effect (**Fig. 3C**). Silencing of *NFU1* significantly lowered ^55^Fe incorporation to ∼10% of that in non-silenced cells. As a control, we also examined ^55^Fe incorporation in the *bS16m*-KO cell line reconstituted with the bS16mC26A variant that supports ∼60% of WT mitochondrial protein synthesis (**Fig. 1H-I**), wherein we detected ∼25% of ^55^Fe as compared to the WT cell line.

To determine the consequences of attenuating [Fe-S] cluster delivery proteins on mitoribosome function, we analyzed the incorporation of ^35^S-methionine into mitoribosome-synthesized polypeptides in the presence of emetine to inhibit cytosolic protein synthesis. Consistent with the mitoribosome stability and ^55^Fe-incorporation assays, the silencing of GLRX5, ISCA1, BOLA3, and NFU1, but not NUBPL, diminished the mitochondrial protein synthesis capacity to 60-70 % of control (**Fig. 3D**).

### Mutation in multiple mitochondrial dysfunctions syndrome (MMDS) genes *BOLA3* and *NFU1* disrupt mitoribosome assembly and function

Mutations in *NFU1* or *BOLA3* result in MMDS types 1 and 2, respectively (33). We have shown in a previous study that fibroblasts from subjects carrying mutations in either *NFU1* or *BOLA3* showed defective delivery of [Fe-S] clusters to OXPHOS complexes I, II, and III, as well as attenuated lipoate synthesis (31). These phenotypes were rescued by the overexpression of mitochondrial isoforms mitoNFU1 (Q9UMS0-1) and BOLA3.1 (Q53S33-1), respectively, but not the cytosolic isoforms cytoNFU1 (Q9UMS0-3) and BOLA3.2 (Q53S33-2) (31). Other studies have shown that mutations in NFU1 also cause dysregulation of cellular iron as well as oxidative stress in patient cell lines and model organisms (57,58), which may further affect the integrity of redox-sensitive mitochondrial [Fe-S] clusters, such as those in mS25-bS16m and METTL17 (**Fig. 2C-H**)

Here, we have analyzed the steady-state levels of mitoribosome proteins in subject fibroblasts to determine the effect of NFU1 and BOLA3 variants on mitoribosome maturation. (**Fig. 4A**). In contrast to siRNA-mediated knockdown of NFU1 or BOLA3 (**Supplementary Fig. S5E**), mitoribosome protein levels were not decreased in subject fibroblasts (**Fig. 4A**). However, overexpression of mitoNFU1 and BOLA3.1 resulted in increases of several mitoribosome markers, particularly those involved in [2Fe-2S] coordination (bS16m, bS18m, mS25, and uL10m). NFU1 levels were not altered in BOLA3 subject fibroblasts. Interestingly, NFU1 subject fibroblasts presented with decreased levels of both NFU1 and BOLA3 proteins, an effect not seen in siRNA-mediated knock-down. As shown previously (31), the levels of subunits from OXPHOS complexes containing [Fe-S] clusters were significantly decreased in subject fibroblasts and rescued by mitoNFU1 and BOLA3.1, respectively. Additionally, the steady-state levels of Complex IV subunits were also lowered, probably reflecting the dependency of Complex IV assembly on [Fe-S] cluster-containing proteins (16,59). Decreased Complex IV levels could be partly attributed to potentially defective mitochondrial translation in subject fibroblasts, either due to inefficient protein synthesis or defective mitoribosome assembly. In support of this possibility, an *in cellula* translation assay showed a clear defect in the pulse synthesis of mtDNA-encoded proteins (**Fig. 4B-C**), and sucrose gradient sedimentation analyses showed decreased mtSSU and monosome complexes in NFU1 subject fibroblasts (**Fig. 4D**) in agreement with the role of NFU1 in delivering a [4Fe-4S] cluster to the mtSSU-specific assembly factor METTL17. The mitochondrial protein synthesis defect resulted in deficient OXPHOS complex assembly (**Fig. 4E**). All presented phenotypes were rescued by overexpression of mitoNFU1 and BOLA3.1, respectively. In summary, the results in subject fibroblasts largely phenocopy our findings in siRNA-mediated *NFU1* and *BOLA3* knockdown cells and support the conclusion that attenuated mitoribosome assembly and function contribute to the molecular phenotype of MMDS types 1 and 2.

**Figure 4.**
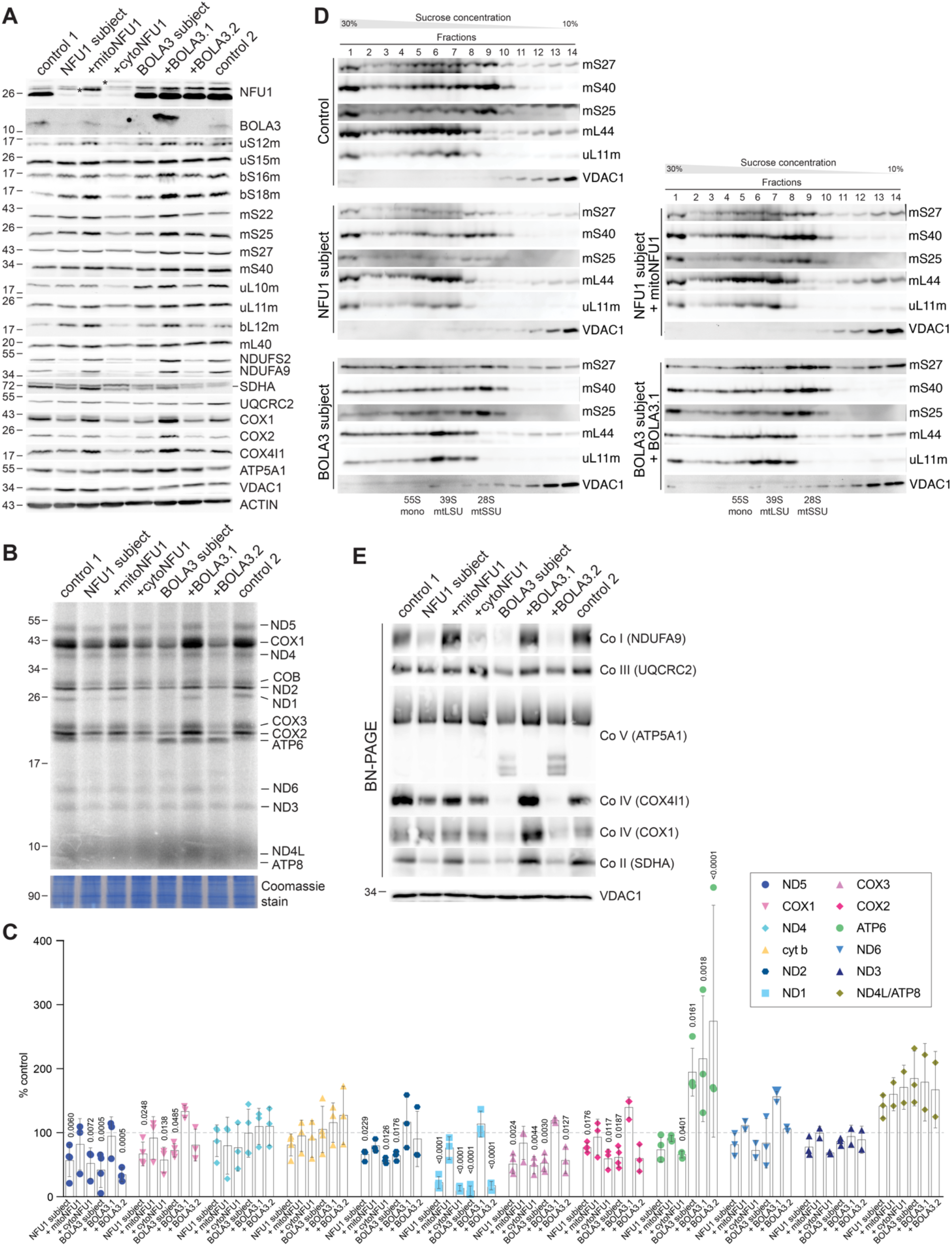
Multiple mitochondrial dysfunctions syndrome-associated proteins BOLA3 and NFU1 are important for mitoribosome assembly and function. (**A**) Steady-state levels of indicated mitoribosomal proteins and subunits of OXPHOS complexes were assessed by immunoblotting in fibroblasts from two unrelated controls, subjects carrying mutations in either *NFU1* or *BOLA3*, or subjects overexpressing the rescuing mitochondrial isoforms mitoNFU1 (Q9UMS0-1) and BOLA3.1 (Q53S33-1), or non-rescuing cytosolic isoforms cytoNFU1 (Q9UMS0-3) and BOLA3.2 (Q53S33-2). VDAC1 and ACTIN were used as loading controls. (**B**) Impaired mitochondrial protein synthesis in NFU1 and BOLA3 subject fibroblasts following ^35^S-methionine incorporation into newly synthesized mitochondrial polypeptides in the presence of emetine to inhibit cytoplasmic protein synthesis. A Coomassie total protein staining served as a loading control. (**C**) Quantification of at least three independent experiments as in panel B. The graphs show the densitometry (average ± SD) for each indicated polypeptide. Shapes represent individual data points. Two-way ANOVA with a Dunnett correction for multiple comparisons was performed (compared to controls, which were set to 100%), and the significant *p*-values are indicated. (**D**) Sucrose gradient sedimentation analyses of mtSSU (mS27, mS40, and mS25) and mtLSU markers (mL44, uL11m) from mitochondria purified from the indicated cell lines. VDAC1 was used as a loading control. (**E**) BN-PAGE analysis of the assembly of OXPHOS complexes in indicated cell lines using subunit-specific antibodies (indicated in parenthesis). BN-PAGE samples were separated on an SDS-PAGE, and VDAC1 was used as a loading control.

## Discussion

Iron-sulfur clusters are critical co-factors that function in electron transfer and storage, chemical catalysis, structural stabilization, and sensing oxygen radicals and other small molecules (60,61). The recent discovery of three [2Fe-2S] clusters in the mitoribosome and one [4Fe-4S] cluster in a mtSSU assembly factor has linked the synthesis of OXPHOS core subunits with their assembly into [Fe-S]-containing enzymatic units. Furthermore, diseases associated with defective iron-sulfur cluster biosynthesis stem not only from altered OXPHOS complex assembly, but also from impaired synthesis of mtDNA-encoded subunits. In this study, we have unraveled the role of the mitoribosome [2Fe-2S] clusters in the structural stabilization of the mtSSU and oxidative stress sensing to regulate mitochondrial translation accordingly. By combining genetic silencing of ISC machinery and quantitative proteomic analyses, we identified GLRX5 and BOLA3 as the mitoribosome [2Fe-2S] cluster delivery factors and ISCA1 and NFU1 as factors that target a [4Fe-4S] cluster to the mtSSU assembly protein METTL17 (**Fig. 3E**). Finally, we used fibroblasts from subjects with mutations in NFU1 and BOLA3 to disclose previously unrecognized mitochondrial protein synthesis defects that may contribute to pathophysiology of these mutations. In this fashion, our data link mitoribosomal biogenesis, Fe-S cluster assembly, and the pathophysiology of subjects suffering from “multiple mitochondrial dysfunction” syndromes.

A functional link between cytosolic ribosome assembly and biogenesis of iron-sulfur proteins was previously established in the yeast *Saccharomyces cerevisiae*, wherein the [4Fe-4S] protein RNase L inhibitor (Rli1) functions in several ribosome assembly steps (62,63), the process of translation (64,65), and ribosome recycling (66). The [4Fe-4S] iron-sulfur cluster in Rli1 senses reactive oxygen species (ROS) and – through a loss of activity – attenuates protein synthesis under oxidative stress conditions (67). In *Escherichia coli,* a [4Fe-4S] iron-sulfur cluster was also found in RumA, the enzyme that methylates U1939 23S rRNA (68). In this case, the cluster also acts as a ROS sensor. Oxidation of said cluster leads to its decomposition and subsequent precipitation of the protein, which may constitute a mechanism for regulating RumA activity upon oxidative insults (69). The absence of RumA-mediated 23S rRNA methylation might be advantageous to ribosomal functions under oxidative stress conditions. Another interesting hypothesis proposed for RumA regulation is that because far fewer turnovers of an enzyme involved in rRNA modification are required than of other enzymes and electron transfer [Fe-S] cluster-containing proteins, the cluster in RumA may be a sacrificial one to serve as iron and sulfur donor for other clusters after the process of 23S rRNA methylation is completed (69). Our data show that in the mitoribosome, at least the [2Fe-2S] cluster coordinated by mS25-bS16m is sensitive to ROS – likely to attenuate mitochondrial protein synthesis under oxidative stress conditions readily. Additionally, the [4Fe-4S] cluster in the methyltransferase-like mitoribosome assembly factor METTL17, required for mtSSU assembly, is highly sensitive to H_2_O_2_, likely to prevent new mtSSU assembly under oxidative insults. In that regard, METTL17 acts similarly to the yeast cytosolic and bacterial ribosome assembly factors mentioned earlier. Although METTL17 does not seem to possess methyltransferase activity, it associates with the 12S rRNA in the mtSSU head to promote the rearrangement of several helices and block the mRNA channel to prevent its premature binding (8). The role of METTL17 seems to be conserved along evolution. A cryo-EM reconstruction showed that both its yeast and trypanosomid homologs, known as Rsm22, also interact with and contribute to the maturation of mitoribosome mtSSU assembly intermediates (56,70). However, whereas yeast Rsm22 was found to coordinate a [4Fe-4S] cluster (56), a Zn^2+^ was assigned to the density detected in its trypanosomid counterpart (70), perhaps due to a lack of resolution. Independently, it is well possible that, Rsm22 has the capacity to sense oxidative stress like METTL17 does.

Deficiencies in the majority of [Fe-S] cluster biogenesis and assembly factors have been associated with an array of human mitochondrial and neurodegenerative disorders, including Friedrich ataxia (71) and multiple mitochondrial dysfunctions syndrome (33). A common hallmark of these disorders is mitochondrial oxidative phosphorylation deficiency observed in subject samples (33,71). This is expected as respiratory complexes I, II and III contain [Fe-S] clusters and the biogenesis of Complex IV depends on [Fe-S] cluster-containing proteins (16,59). However, the contribution of mitoribosomal depletion has not been previously considered. Our data show that fibroblasts from subjects carrying variants in [Fe-S] cluster biogenesis factors BOLA3 or NFU1 exhibit attenuated mitochondrial protein synthesis, which cannot be accounted for by defective-OXPHOS feedback loop and is consistent with the role of these proteins in the delivery of [Fe-S] clusters to the mitoribosome and METTL17, respectively. Therefore, our data adds a new important wrinkle to understanding the pathogenic mechanisms underlying [Fe-S] cluster biogenesis-related deficiencies.

## DATA AND CODE AVAILABILITY

All unique/stable reagents generated in this study (plasmids and cell lines) are available from the Lead Contact with a completed Materials Transfer Agreement.

All Source Data is either included in the manuscript or will be provided upon request. Multiplex TMT-MS/MS data have been uploaded to PRIDE repository, with ref# 1-20230106-165123.

## SUPPLEMENTARY DATA

Figures S1-S5, and Table S1-S2

## ACKNOWLEDGEMENTS

We thank Dr. F. Fontanesi for the critical reading of the manuscript. We thank the Bioinformatics and Statistics Core at the University of Florida-Scripps Biomedical Research for MS data generation and analysis. We thank Kathleen Daigneault for her technical assistance.

## Author contributions

HZ and AB performed biochemical characterization and cell biology studies. HZ created and characterized the HEK293T KO lines and performed most experiments with HEK293T. AB contributed with OXPHOS measurements. AJ, HA, and ES designed and performed the experiments with *NFU1* and *BOLA3* subject fibroblasts. OK provided intellectual input in the experimental design for experiments on redox sensing and ^55^Fe incorporation. HZ and AB wrote the first draft of the paper, and AJ, HA, and ES added the results on subject fibroblasts. All authors contributed to data interpretation and read, edited, and approved the manuscript.

## REFERENCES

1. Kummer, E. and Ban, N. (2021) Mechanisms and regulation of protein synthesis in mitochondria. Nat. Rev. Mol. Cell Biol., 22, 307–325.

2. Ferrari, A., Del’Olio, S. and Barrientos, A. (2021) The diseased mitoribosome. FEBS Lett, 595, 1025–1061.

3. Kim, H.J., Maiti, P. and Barrientos, A. (2017) Mitochondrial ribosomes in cancer. Semin. Cancer Biol., 47, 67–81.

4. Itoh, Y., Khawaja, A., Laptev, I., Cipullo, M., Atanassov, I., Sergiev, P., Rorbach, J. and Amunts, A. (2022) Mechanism of mitoribosomal small subunit biogenesis and preinitiation. Nature, 606, 603–608.

5. Tobiasson, V., Gahura, O., Aibara, S., Baradaran, R., Zíková, A. and Amunts, A. (2021) Interconnected assembly factors regulate the biogenesis of mitoribosomal large subunit. Embo J., 40, e106292.

6. Bugiardini, E., Mitchell, A.L., Rosa, I.D., Horning-Do, H.T., Pitmann, A., Poole, O.V., Holton, J.L., Shah, S., Woodward, C., Hargreaves, I. et al. (2019) MRPS25 mutations impair mitochondrial translation and cause encephalomyopathy. Hum. Mol. Genet., 30.

7. Miller, C., Saada, A., Shaul, N., Shabtai, N., Ben-Shalom, E., Shaag, A., Hershkovitz, E. and Elpeleg, O. (2004) Defective mitochondrial translation caused by a ribosomal protein (MRPS16) mutation. Ann. Neurol., 56, 734–738.

8. Ast, T., Itoh, Y., Sadre, S., McCoy, J.G., Namkoong, G., Chicherin, I.V., Joshi, P.R., Kamenski, P., Suess, D.L.M., Amunts, A. et al. (2022) METTL17 is an Fe-S cluster checkpoint for mitochondrial translation. bioRxiv, 2022.2011.2024.517765.

9. Camponeschi, F., Ciofi-Baffoni, S., Calderone, V. and Banci, L. (2022) Molecular Basis of Rare Diseases Associated to the Maturation of Mitochondrial [4Fe-4S]-Containing Proteins. Biomolecules, 12.

10. Maio, N. and Rouault, T.A. (2020) Outlining the Complex Pathway of Mammalian Fe-S Cluster Biogenesis. Trends Biochem. Sci., 45, 411–426.

11. Lill, R. and Freibert, S.A. (2020) Mechanisms of Mitochondrial Iron-Sulfur Protein Biogenesis. Annu. Rev. Biochem., 89, 471–499.

12. Braymer, J.J., Freibert, S.A., Rakwalska-Bange, M. and Lill, R. (2021) Mechanistic concepts of iron-sulfur protein biogenesis in Biology. Biochim. Biophys. Acta Mol. Cell Res., 1868, 118863.

13. Boniecki, M.T., Freibert, S.A., Mühlenhoff, U., Lill, R. and Cygler, M. (2017) Structure and functional dynamics of the mitochondrial Fe/S cluster synthesis complex. Nature communications, 8, 1287.

14. Van Vranken, J.G., Jeong, M.Y., Wei, P., Chen, Y.C., Gygi, S.P., Winge, D.R. and Rutter, J. (2016) The mitochondrial acyl carrier protein (ACP) coordinates mitochondrial fatty acid synthesis with iron sulfur cluster biogenesis. Elife, 5.

15. Cory, S.A., Van Vranken, J.G., Brignole, E.J., Patra, S., Winge, D.R., Drennan, C.L., Rutter, J. and Barondeau, D.P. (2017) Structure of human Fe-S assembly subcomplex reveals unexpected cysteine desulfurase architecture and acyl-ACP-ISD11 interactions. Proc. Natl. Acad. Sci. U. S. A., 114, E5325–e5334.

16. Sheftel, A.D., Stehling, O., Pierik, A.J., Elsässer, H.P., Mühlenhoff, U., Webert, H., Hobler, A., Hannemann, F., Bernhardt, R. and Lill, R. (2010) Humans possess two mitochondrial ferredoxins, Fdx1 and Fdx2, with distinct roles in steroidogenesis, heme, and Fe/S cluster biosynthesis. Proc. Natl. Acad. Sci. U. S. A., 107, 11775–11780.

17. Cai, K., Tonelli, M., Frederick, R.O. and Markley, J.L. (2017) Human Mitochondrial Ferredoxin 1 (FDX1) and Ferredoxin 2 (FDX2) Both Bind Cysteine Desulfurase and Donate Electrons for Iron-Sulfur Cluster Biosynthesis. Biochemistry, 56, 487–499.

18. Dutkiewicz, R. and Nowak, M. (2018) Molecular chaperones involved in mitochondrial iron-sulfur protein biogenesis. J. Biol. Inorg. Chem., 23, 569–579.

19. Nasta, V., Giachetti, A., Ciofi-Baffoni, S. and Banci, L. (2017) Structural insights into the molecular function of human [2Fe-2S] BOLA1-GRX5 and [2Fe-2S] BOLA3-GRX5 complexes. Biochim. Biophys. Acta Gen. Subj., 1861, 2119–2131.

20. Uzarska, M.A., Nasta, V., Weiler, B.D., Spantgar, F., Ciofi-Baffoni, S., Saviello, M.R., Gonnelli, L., Mühlenhoff, U., Banci, L. and Lill, R. (2016) Mitochondrial Bol1 and Bol3 function as assembly factors for specific iron-sulfur proteins. Elife, 5.

21. Nasta, V., Suraci, D., Gourdoupis, S., Ciofi-Baffoni, S. and Banci, L. (2020) A pathway for assembling [4Fe-4S](2+) clusters in mitochondrial iron-sulfur protein biogenesis. Febs J., 287, 2312–2327.

22. Braymer, J.J. and Lill, R. (2017) Iron-sulfur cluster biogenesis and trafficking in mitochondria. J. Biol. Chem., 292, 12754–12763.

23. Maio, N. and Rouault, T.A. (2022) Mammalian iron sulfur cluster biogenesis: From assembly to delivery to recipient proteins with a focus on novel targets of the chaperone and co-chaperone proteins. IUBMB Life, 74, 684–704.

24. Bych, K., Kerscher, S., Netz, D.J., Pierik, A.J., Zwicker, K., Huynen, M.A., Lill, R., Brandt, U. and Balk, J. (2008) The iron-sulphur protein Ind1 is required for effective complex I assembly. EMBO J., 27, 1736–1746.

25. Melber, A., Na, U., Vashisht, A., Weiler, B.D., Lill, R., Wohlschlegel, J.A. and Winge, D.R. (2016) Role of Nfu1 and Bol3 in iron-sulfur cluster transfer to mitochondrial clients. Elife, 5.

26. Rötig, A., de Lonlay, P., Chretien, D., Foury, F., Koenig, M., Sidi, D., Munnich, A. and Rustin, P. (1997) Aconitase and mitochondrial iron-sulphur protein deficiency in Friedreich ataxia. Nat. Genet., 17, 215–217.

27. Mochel, F., Knight, M.A., Tong, W.H., Hernandez, D., Ayyad, K., Taivassalo, T., Andersen, P.M., Singleton, A., Rouault, T.A., Fischbeck, K.H. et al. (2008) Splice mutation in the iron-sulfur cluster scaffold protein ISCU causes myopathy with exercise intolerance. Am. J. Hum. Genet., 82, 652–660.

28. Olsson, A., Lind, L., Thornell, L.E. and Holmberg, M. (2008) Myopathy with lactic acidosis is linked to chromosome 12q23.3-24.11 and caused by an intron mutation in the ISCU gene resulting in a splicing defect. Hum. Mol. Genet., 17, 1666–1672.

29. Camaschella, C., Campanella, A., De Falco, L., Boschetto, L., Merlini, R., Silvestri, L., Levi, S. and Iolascon, A. (2007) The human counterpart of zebrafish shiraz shows sideroblastic-like microcytic anemia and iron overload. Blood, 110, 1353–1358.

30. Feng, W.X., Zhuo, X.W., Liu, Z.M., Li, J.W., Zhang, W.H., Wu, Y., Han, T.L. and Fang, F. (2021) Case Report: A variant non-ketotic hyperglycinemia with GLRX5 mutations: manifestation of deficiency of activities of the respiratory chain enzymes. Front. Genet., 12, 605778.

31. Cameron, J.M., Janer, A., Levandovskiy, V., Mackay, N., Rouault, T.A., Tong, W.H., Ogilvie, I., Shoubridge, E.A. and Robinson, B.H. (2011) Mutations in iron-sulfur cluster scaffold genes NFU1 and BOLA3 cause a fatal deficiency of multiple respiratory chain and 2-oxoacid dehydrogenase enzymes. Am. J. Hum. Genet., 89, 486–495.

32. Navarro-Sastre, A., Tort, F., Stehling, O., Uzarska, M.A., Arranz, J.A., Del Toro, M., Labayru, M.T., Landa, J., Font, A., Garcia-Villoria, J. et al. (2011) A fatal mitochondrial disease is associated with defective NFU1 function in the maturation of a subset of mitochondrial Fe-S proteins. Am. J. Hum. Genet., 89, 656–667.

33. Lebigot, E., Schiff, M. and Golinelli-Cohen, M.P. (2021) A review of Multiple Mitochondrial Dysfunction Syndromes, syndromes associated with defective Fe-S protein maturation. Biomedicines, 9.

34. Morita, E., Arii, J., Christensen, D., Votteler, J. and Sundquist, W.I. (2012) Attenuated protein expression vectors for use in siRNA rescue experiments. Biotechniques, 0, 1–5.

35. Bourens, M., Boulet, A., Leary, S.C. and Barrientos, A. (2014) Human COX20 cooperates with SCO1 and SCO2 to mature COX2 and promote the assembly of cytochrome *c* oxidase. Hum. Mol. Genet., 23, 2901–2913.

36. Choi, A. and Barrientos, A. (2021) Sucrose gradient sedimentation analysis of mitochondrial ribosomes. Methods Mol. Biol., 2192, 211–226.

37. Schindelin, J., Arganda-Carreras, I., Frise, E., Kaynig, V., Longair, M., Pietzsch, T., Preibisch, S., Rueden, C., Saalfeld, S., Schmid, B. et al. (2012) Fiji: an open-source platform for biological-image analysis. Nat. Methods, 9, 676–682.

38. Barrientos, A., Fontanesi, F. and Diaz, F. (2009) Evaluation of the mitochondrial respiratory chain and oxidative phosphorylation system using polarography and spectrophotometric enzyme assays. Curr. Protoc. Hum. Genet., Chapter, Unit19.13.

39. Lowry, O.H., Rosebrough, N.J., Farr, A.L. and Randall, R.J. (1951) Protein measurement with the Folin phenol reagent. J. Biol. Chem., 193, 265–275.

40. Bradford, M.M. (1976) A rapid and sensitive method for the quantitation of microgram quantities of protein utilizing the principle of protein-dye binding. Anal. Biochem., 72, 248–254.

41. Laemmli, U.K. (1970) Cleavage of structural proteins during the assembly of the head of bacteriophage T4. Nature, 227, 680–685.

42. Nývltová, E., Dietz, J.V., Seravalli, J., Khalimonchuk, O. and Barrientos, A. (2022) Coordination of metal center biogenesis in human cytochrome *c* oxidase. Nature communications, 13, 3615.

43. Soto, I.C. and Barrientos, A. (2016) Mitochondrial cytochrome *c* oxidase biogenesis is regulated by the redox state of a heme-binding translational activator. Antioxid. Redox Signal., 24, 281–298.

44. Maio, N., Singh, A., Uhrigshardt, H., Saxena, N., Tong, W.H. and Rouault, T.A. (2014) Cochaperone binding to LYR motifs confers specificity of iron sulfur cluster delivery. Cell Metab., 19, 445–457.

45. Antonicka, H., Ogilvie, I., Taivassalo, T., Anitori, R.P., Haller, R.G., Vissing, J., Kennaway, N.G. and Shoubridge, E.A. (2003) Identification and characterization of a common set of complex I assembly intermediates in mitochondria from patients with complex I deficiency. J. Biol. Chem., 278, 43081–43088.

46. Leary, S.C. and Sasarman, F. (2009) Oxidative phosphorylation: synthesis of mitochondrially encoded proteins and assembly of individual structural subunits into functional holoenzyme complexes. Methods Mol. Biol., 554, 143–162.

47. Koh, C.M. (2013) Isolation of genomic DNA from mammalian cells. Methods Enzymol., 529, 161–169.

48. Bustin, S.A., Benes, V., Garson, J.A., Hellemans, J., Huggett, J., Kubista, M., Mueller, R., Nolan, T., Pfaffl, M.W., Shipley, G.L. et al. (2009) The MIQE guidelines: minimum information for publication of quantitative real-time PCR experiments. Clin. Chem., 55, 611–622.

49. Lefever, S., Hellemans, J., Pattyn, F., Przybylski, D.R., Taylor, C., Geurts, R., Untergasser, A. and Vandesompele, J. (2009) RDML: structured language and reporting guidelines for real-time quantitative PCR data. Nucleic Acids Res., 37, 2065–2069.

50. Pfaffl, M.W. (2001) A new mathematical model for relative quantification in real-time RT-PCR. Nucleic Acids Res., 29, e45.

51. Käll, L., Storey, J.D. and Noble, W.S. (2008) Non-parametric estimation of posterior error probabilities associated with peptides identified by tandem mass spectrometry. Bioinformatics, 24, i42–48.

52. Nesvizhskii, A.I., Keller, A., Kolker, E. and Aebersold, R. (2003) A statistical model for identifying proteins by tandem mass spectrometry. Anal. Chem., 75, 4646–4658.

53. Oberg, A.L., Mahoney, D.W., Eckel-Passow, J.E., Malone, C.J., Wolfinger, R.D., Hill, E.G., Cooper, L.T., Onuma, O.K., Spiro, C., Therneau, T.M. et al. (2008) Statistical analysis of relative labeled mass spectrometry data from complex samples using ANOVA. J. Proteome Res., 7, 225–233.

54. Crack, J.C. and Le Brun, N.E. (2018) Redox-sensing iron-sulfur cluster regulators. Antioxid. Redox Signal., 29, 1809–1829.

55. Ransy, C., Vaz, C., Lombès, A. and Bouillaud, F. (2020) Use of H(2)O(2) to cause oxidative stress, the catalase issue. Int. J. Mol. Sci., 21.

56. Harper, N.J., Burnside, C. and Klinge, S. (2023) Principles of mitoribosomal small subunit assembly in eukaryotes. Nature, 614, 175–181.

57. Wachnowsky, C., Wesley, N.A., Fidai, I. and Cowan, J.A. (2017) Understanding the molecular basis of Multiple Mitochondrial Dysfunctions Syndrome 1 (MMDS1)-impact of a dsease-causing Gly208Cys substitution on structure and activity of NFU1 in the Fe/S cluster biosynthetic pathway. J. Mol. Biol., 429, 790–807.

58. Kropp, P.A., Wu, J., Reidy, M., Shrestha, S., Rhodehouse, K., Rogers, P., Sack, M.N. and Golden, A. (2021) Allele-specific mitochondrial stress induced by Multiple Mitochondrial Dysfunctions Syndrome 1 pathogenic mutations modeled in *Caenorhabditis elegans*. PLoS Genet., 17, e1009771.

59. Lange, H., Mühlenhoff, U., Denzel, M., Kispal, G. and Lill, R. (2004) The heme synthesis defect of mutants impaired in mitochondrial iron-sulfur protein biogenesis is caused by reversible inhibition of ferrochelatase. J. Biol. Chem., 279, 29101–29108.

60. Schulz, V., Freibert, S.A., Boss, L., Mühlenhoff, U., Stehling, O. and Lill, R. (2023) Mitochondrial [2Fe-2S] ferredoxins: new functions for old dogs. FEBS Lett., 597, 102–121.

61. Rouault, T.A. (2019) The indispensable role of mammalian iron sulfur proteins in function and regulation of multiple diverse metabolic pathways. Biometals, 32, 343–353.

62. Yarunin, A., Panse, V.G., Petfalski, E., Dez, C., Tollervey, D. and Hurt, E.C. (2005) Functional link between ribosome formation and biogenesis of iron-sulfur proteins. EMBO. J., 24, 580–588.

63. Kispal, G., Sipos, K., Lange, H., Fekete, Z., Bedekovics, T., Janáky, T., Bassler, J., Aguilar Netz, D.J., Balk, J., Rotte, C. et al. (2005) Biogenesis of cytosolic ribosomes requires the essential iron-sulphur protein Rli1p and mitochondria. EMBO J., 24, 589–598.

64. Khoshnevis, S., Gross, T., Rotte, C., Baierlein, C., Ficner, R. and Krebber, H. (2010) The iron-sulphur protein RNase L inhibitor functions in translation termination. EMBO Rep., 11, 214–219.

65. Dong, J., Lai, R., Nielsen, K., Fekete, C.A., Qiu, H. and Hinnebusch, A.G. (2004) The essential ATP-binding cassette protein RLI1 functions in translation by promoting preinitiation complex assembly. J. Biol. Chem., 279, 42157–42168.

66. Barthelme, D., Dinkelaker, S., Albers, S.V., Londei, P., Ermler, U. and Tampé, R. (2011) Ribosome recycling depends on a mechanistic link between the FeS cluster domain and a conformational switch of the twin-ATPase ABCE1. Proc. Natl. Acad. Sci. U. S. A., 108, 3228–3233.

67. Alhebshi, A., Sideri, T.C., Holland, S.L. and Avery, S.V. (2012) The essential iron-sulfur protein Rli1 is an important target accounting for inhibition of cell growth by reactive oxygen species. Mol. Biol. Cell, 23, 3582–3590.

68. Lee, T.T., Agarwalla, S. and Stroud, R.M. (2004) Crystal structure of RumA, an iron-sulfur cluster containing E. coli ribosomal RNA 5-methyluridine methyltransferase. Structure, 12, 397–407.

69. Agarwalla, S., Stroud, R.M. and Gaffney, B.J. (2004) Redox reactions of the iron-sulfur cluster in a ribosomal RNA methyltransferase, RumA: optical and EPR studies. J. Biol. Chem., 279, 34123–34129.

70. Saurer, M., Ramrath, D.J.F., Niemann, M., Calderaro, S., Prange, C., Mattei, S., Scaiola, A., Leitner, A., Bieri, P., Horn, E.K. et al. (2019) Mitoribosomal small subunit biogenesis in trypanosomes involves an extensive assembly machinery. Science, 365, 1144–1149.

71. Rotig, A., de Lonlay, P., Chretien, D., Foury, F., Koenig, M., Sidi, D., Munnich, A. and Rustin, P. (1997) Aconitase and mitochondrial iron-sulphur protein deficiency in Friedreich ataxia. Nat. Genet., 17, 215–217.

