## Supplementary Data for "BOLA3 and NFU1 link mitoribosome iron-sulfur cluster assembly to multiple mitochondrial dysfunctions syndrome"

**This PDF file includes:**

Supplementary Figures 1 to 5
Supplementary Table 1 to 2

### Supplementary Figures

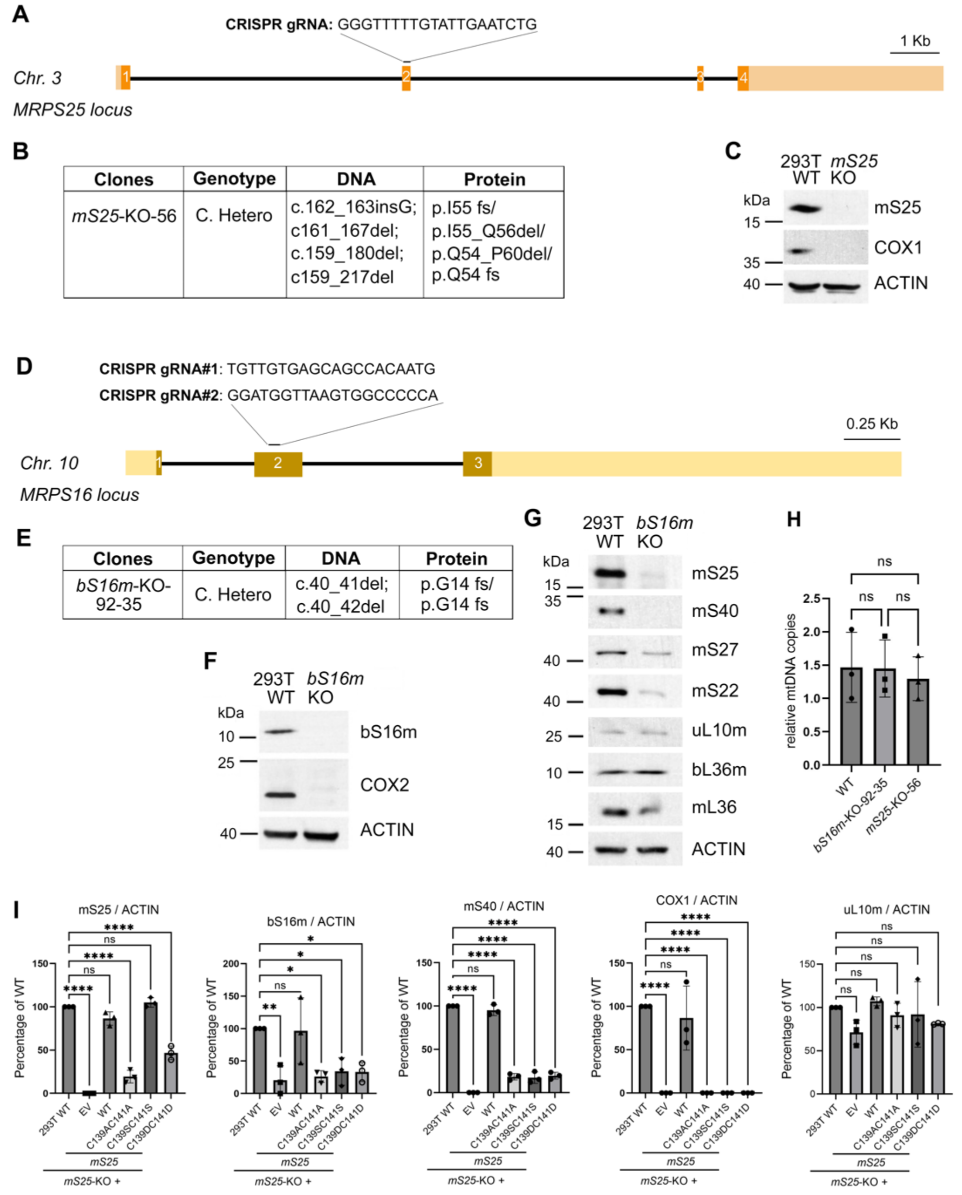

**Figure S1. Generation of *mS25*-KO, and *bS16m*-KO cell lines in the HEK293T background.** (**A** and **D**) Schematic showing the location of the target sites in the *mS25*, and *bS16m* loci for the CRISPR guide RNAs used to knock out each gene. *Chr.*, chromosome. (**B** and **E**) Table detailing the genotype of the generated HEK293T cell lines carrying edited *mS25*, or *bS16m* alleles. *C. Hetero*, compound heterozygous. (**C, F,** and **G**) Immunoblot validation of the respective KO clones. Once the KO cell lines for each relevant gene were established, each was tested three more times, and frozen stocks were stored in liquid nitrogen. Every time a new frozen vial was thawed, the KO cell lines were re-validated by immunoblotting. (**H**) qPCR estimation of mtDNA copy number in HEK293T WT, *bS16m*-KO-92-35, and *mS25*-KO-56. The results were normalized to WT values. Data are presented as mean ± SEM. Statistical differences between the two groups were examined by one-way analysis of variance (ANOVA) followed by Tukey's post hoc test (*p* value<0.05). (**I**) The graphs show the densitometry (average ± SD) of three independent experiments, as in Figure 1B. Black dots represent individual data points. Two-tailed unpaired *t*-test, $^{*}p < 0.05$, $^{**}p < 0.01$, $^{***}p < 0.001$, $^{****}p < 0.0001$.

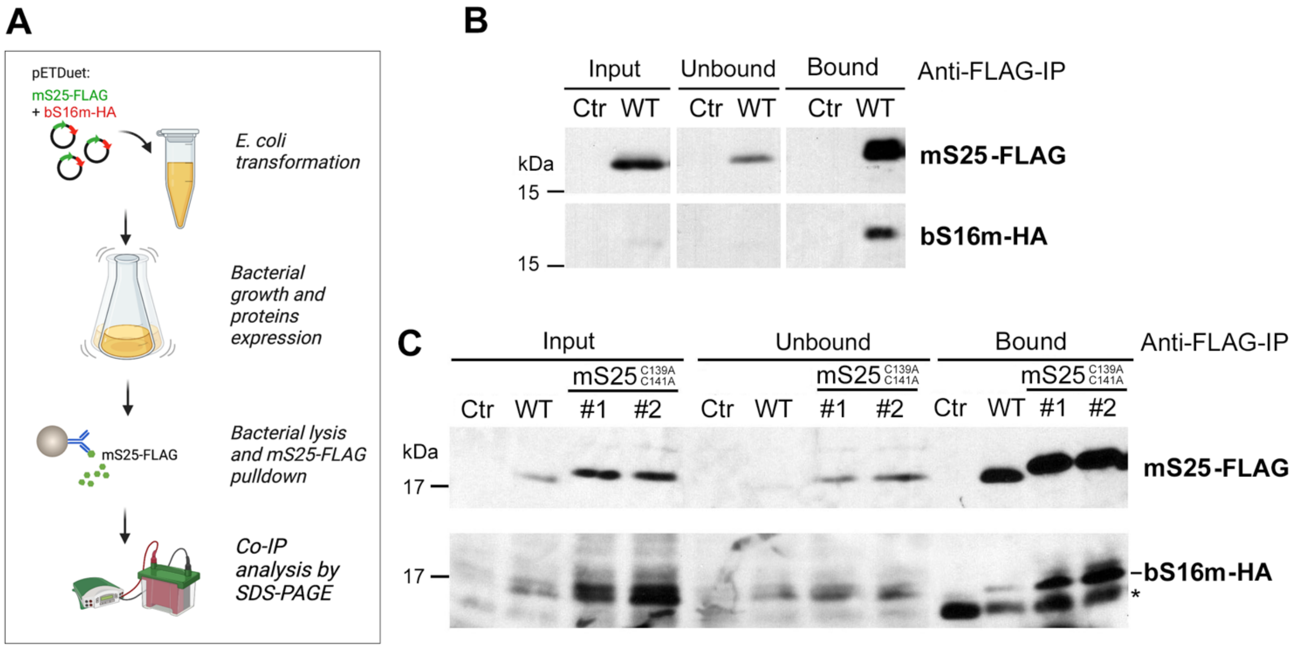

**Figure S2. The proteins mS25 and bS16 interact when co-expressed in *E. coli*.**
(**A**) Schematic representing the experimental approach. mS25-FLAG and bS16m-HA were co-expressed from a pETDuet-1 plasmid in OverExpress C43 (DE3) competent *E. coli* cells, cultured overnight at 30 ºC and lysed. The cleared supernatants were used for immunoprecipitation assays with anti-FLAG-conjugated beads. The corresponding input, unbound, and eluate samples were analyzed by immunoblotting with indicated antibodies.
(**B** and **C**) Co-immunoprecipitation of WT mS25-FLAG (B) or mutant mS25$^{C139AC141A}$-FLAG (C) with bS16m-HA co-expressed in *E. coli* as depicted in panel A. The asterisk denotes unspecific cross-reacting band.

**Figure S3. The steady-state levels of mitoribosome proteins and METTL17 remain changed upon cellular treatment with $H_2O_2$.**
**(A)** Steady-state levels of mS25, bS16m, and METTL17 were assessed by immunoblotting in HEK293T wild-type cells ACTIN was used as a loading control. (**B**) The graphs show the densitometry (average ± SD) of three independent experiments, as in Figure 1B. Black dots represent individual data points. Two-tailed unpaired *t*-test; ns, not-significant.

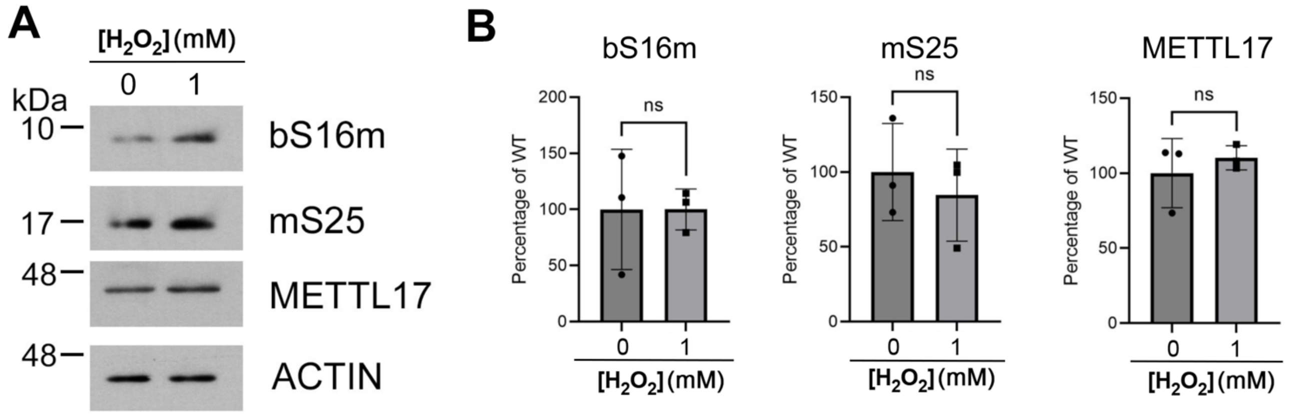

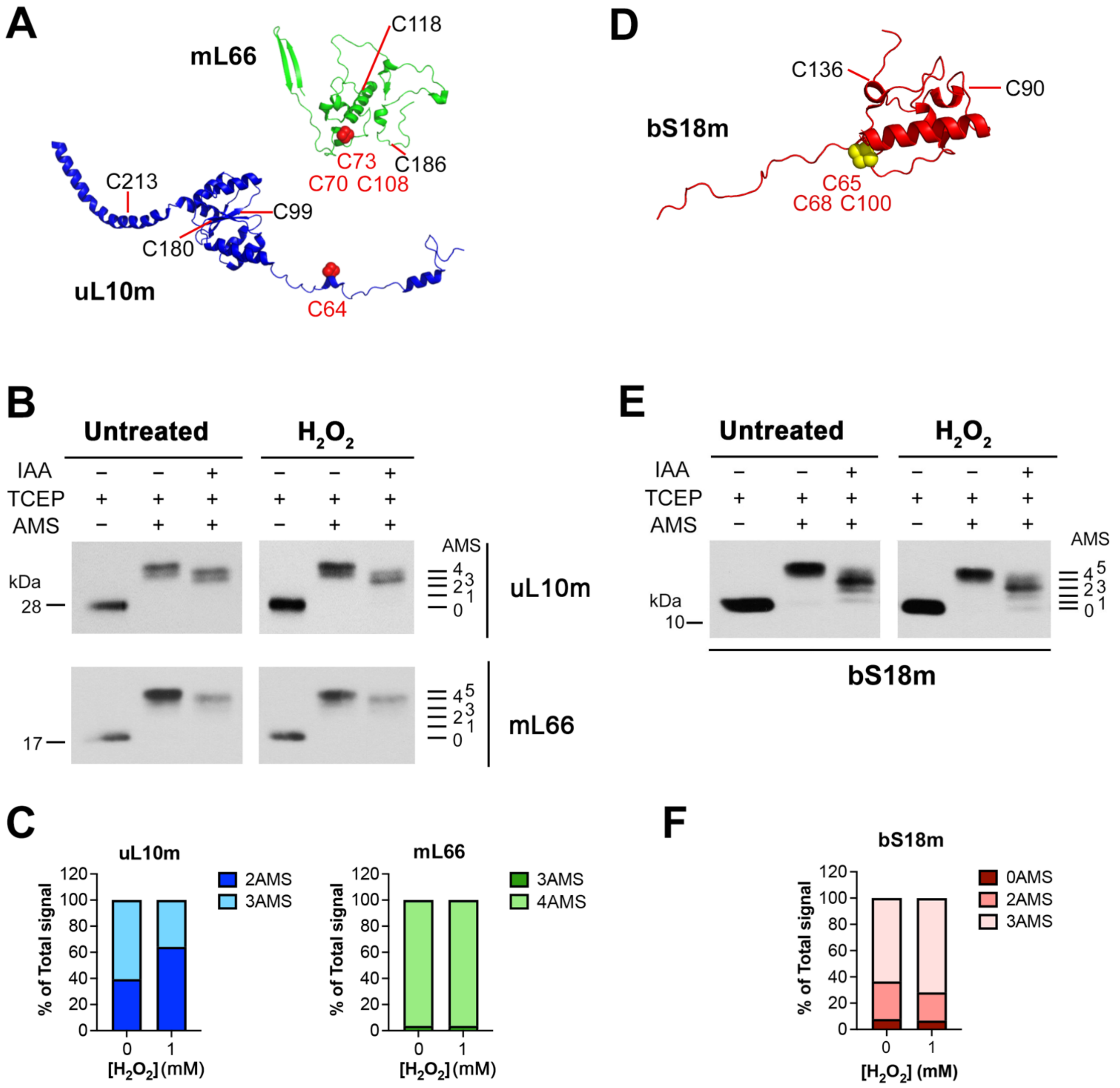

**Figure S4. Cysteines in mL66 and bS18m are not sensitive to oxidative stress.**
(**A** and **D**) Cryo-EM structures (PDB: 7qi4) of the mL66-uL10m pair of proteins that coordinate a [2Fe-2S] cluster in the mtLSU (A), and of bS18m that with bS6m coordinates a [2Fe-2S] in the mtSSU (D). The [2Fe-2S] cluster-coordinating cysteine residues are indicated in red, and all additional cysteines in each protein are shown in black. The [2Fe-2S] cluster is shown (not to scale) as 4 red (A) or yellow (D) spheres. Figures were prepared in Pymol and Adobe Photoshop.
(**B** and **E**) Reverse thiol trapping of mL66, and uL10m (B), and bS18m (E) in mitochondria isolated from WT HEK293T cells, treated or not with 1 mM $H_2O_2$ for 3 h.
(**C** and **F**) The bar graphs represent the number of AMS-bound cysteines (unavailable to IAA binding *in organello*). The data shown are the average of two independent experiments. Figures are representative of two independent repetitions with similar results.

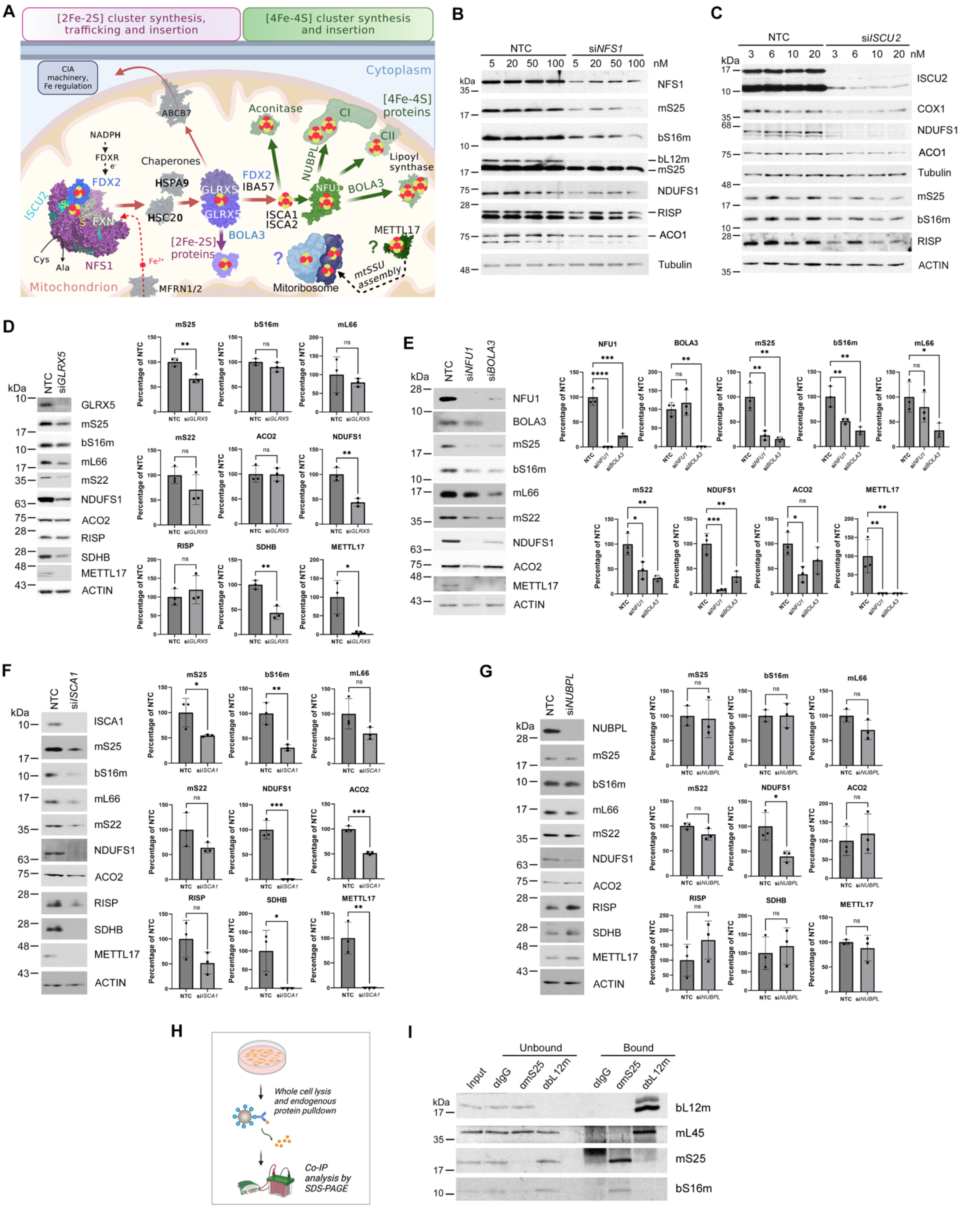

**Figure S5. The lack of [Fe-S] cluster biosynthesis and delivery proteins compromises mitoribosome stability.**

(**A**) Schematic depicting biosynthesis and delivery of Fe-S clusters in mammalian cells. *De novo* Fe-S cluster biosynthesis starts in mitochondria by synthesizing a $[2Fe\text{-}2S]^{2+}$ cluster on a multimeric protein complex, formed by the scaffold protein ISCU2, the cysteine desulfurase complex NFS1/LYRM4/ACP1 that converts Cys into Ala, and frataxin (FXN). Electrons required for the formation of the $[2Fe\text{-}2S]^{2+}$ cluster are provided by the mitochondrial ferredoxin (FDX2)/ferredoxin reductase (FDXR) system. Newly synthesized cluster is then released and transferred - with the help of adaptors such as HSPA9 and HSC20 chaperones - to either a dimer of the monothiol glutaredoxin GLRX5, which passes it to target [2Fe-2S]-binding proteins or to an apo-GLRX5-BOLA3 heterodimeric complex for $[2Fe\text{-}2S]^{2+}$-cluster transfer to yet-to-be-identified targets. The [4Fe-4S] clusters are formed by a specific system that comprises the ISCA1 and ISCA2 proteins and couples two $[2Fe\text{-}2S]^{2+}$ clusters to generate a [4Fe-4S] cluster. The [4Fe-4S] clusters are then transferred to apo-target proteins – directly or through targeting factors. The former is the case for proteins such as mitochondrial aconitase ACO2. In the latter case, a dedicated factor, NFU1, delivers [4Fe-4S] cluster to select apo-target proteins with the assistance of additional factors such as NUBPL for OXPHOS Complex I or BOLA3 for lipoyl synthase. The pathways for [2Fe-2S] delivery to the mitoribosome and [4Fe-4S] delivery to METTL17 remain to be identified.
(**B-G**) Immunoblot analyses of the steady-state levels of NFS1 (B), ISCU (C), GLRX5 (D), and NFU1 and BOLA3 (E), ISCA1 (F), and NUBPL (G) in HEK293T cells after 3 days (panels B, C) or 9 days (panels D-G) transfection with the indicated gene-specific siRNAs. Panels B and C show examples of empirical establishment of the efficient siRNA concentration by titration of si*NFS1* (B) and si*ISCU* (C). Panels D-G show the effect of the determined efficient concentrations of indicated siRNAs. In each case, the steady-state levels of mitoribosomal proteins (mS25, bS16m, mS22, mL66), the [4Fe-4S] cluster-harboring mitoribosome SSU assembly factor METTL17, and mitochondrial proteins containing [2Fe-2S] (NDUFS1 and RISP) or [4Fe-4S] clusters (ACO2, NDUFS1, and SDHB) were also examined. ACTIN or Tubulin served as loading controls. *NTC*, non-targeted control. In panels D-G, the bar graphs on the right-hand side represent densitometric quantification of the steady-state levels of indicated proteins in the images shown on the left. The values were normalized by ACTIN levels and presented as a percentage of siNTC. The bars show the average of three independent replicates ± S.D. Black dots denote individual data points. Two-tailed unpaired *t*-test, ****$p$<0.0001. (**H**) Schematic representing the experimental approach for immunoprecipitation from HEK293T whole-cell extracts. (**I**) Immunoprecipitation of bL12m, mS25, and native interacting MRP proteins from HEK293T WT whole-cell extracts, using antibodies against indicated target proteins and protein A agarose beads. IgG was used as a negative control.

### Supplementary Tables

#### Table S1. TMT mass spectrometry multiplexing strategy.

N=4

| Plex | Label | TMTpro-126 | TMTpro-127N | TMTpro-127C | TMTpro-128N | TMTpro-129C | TMTpro-130N | TMTpro-131C | TMTpro-132N | TMTpro-133C | TMTpro-134N |
|---|---|---|---|---|---|---|---|---|---|---|---|
| 1 | siRNA transfected | NTC | NFU1 | NTC | NFU1 | ISCA1 | GLRX5 | ISCA1 | GLRX5 | BOLA3 | BOLA3 |
| | Repeat | 1 | 1 | 2 | 2 | 1 | 1 | 2 | 2 | 1 | 2 |

| Plex | Label | TMTpro-126 | TMTpro-127N | TMTpro-127C | TMTpro-128N | TMTpro-129C | TMTpro-130N | TMTpro-131C | TMTpro-132N | TMTpro-133C | TMTpro-134N |
|---|---|---|---|---|---|---|---|---|---|---|---|
| 2 | siRNA transfected | ISCA1 | GLRX5 | ISCA1 | GLRX5 | NTC | NFU1 | NTC | NFU1 | BOLA3 | BOLA3 |
| | Repeat | 3 | 3 | 3 | 4 | 3 | 3 | 4 | 4 | 3 | 4 |

#### Table S2. Key resource table.

| REAGENT or RESOURCE | SOURCE | IDENTIFIER |
|---|---|---|
| Antibodies against human proteins | | |
| ATP5A1 | Abcam | Cat# ab14748<br>RRID: AB_301447 |
| β-ACTIN | Abcam | Cat# ab8227<br>RRID: AB_2305186 |
| β-TUBULIN | Sigma | Cat# C4585<br>RRID: AB_258868 |
| BOLA3 | Sigma | Cat# HPA046393<br>Validated enrichment in mitochondrial samples |
| BOLA3 | Sigma | Cat# HPA053162<br>RRID: AB_2682063 |
| ACO1 | Thermo Fisher Scientific | Cat# PA5-82677<br>RRID: AB_2789833 |
| ACO2 | OriGene | Cat# TA500841<br>RRID: AB_11124995 |
| COX1 | Abcam | Cat# ab14705<br>RRID: AB_2084810 |
| COX2/MT-CO2 | Abcam | Cat# ab110258<br>RRID: AB_10887758 |
| COX4I1 | Abcam | Cat# ab110261<br>RRID: AB_10862101 |
| FLAG-tag | Sigma | Cat# F3165<br>RRID: AB_259529 |
| GLRX5 | Sigma | Cat# HPA063716<br>RRID: AB_2685102 |
| GLRX5 | Abnova | Cat# H00051218-M04<br>RRID: AB_10903043 |

| | | |
|---|---|---|
| HSCB | Thermo Fisher Scientific | Cat# PA5-30693<br>RRID: AB_2548167 |
| ISCA1 | Thermo Fisher Scientific | Cat# PA5-60121<br>RRID: AB_2642786 |
| ISCU | Sigma | Cat# HPA057592<br>RRID: AB_2683478 |
| METTL17 | Atlas | Cat# HPA059802<br>RRID: AB_2684129 |
| MRPL10 | Sigma | Cat# HPA021234<br>RRID: AB_1854098 |
| MRPL11 | Cell Signaling Technology | Cat# 2066<br>RRID: AB_2145598 |
| MRPL12 | Abnova | Cat# H00006182-D01P<br>RRID: AB_10642061 |
| MRPL40 | Sigma-Aldrich | Cat# HPA006181<br>RRID: AB_1079411 |
| MRPL44 | ProteinTech | Cat# 16394-1-AP<br>RRID: AB_2146062 |
| MRPS12 | ProteinTech | Cat# 15225-1-AP<br>RRID: AB_10597843 |
| MRPS15 | ProteinTech | Cat# 17006-1-AP<br>RRID: AB_2301068 |
| MRPS16 | Sigma | Cat# HPA050081<br>RRID: AB_2681007 |
| MRPS16 | ProteinTech | Cat# 16735-1-AP<br>RRID: AB_2180166 |
| MRPS18A | Sigma | Cat# HPA035451<br>RRID: AB_10602029 |
| MRPS18B | Sigma | Cat# HPA050334<br>RRID: AB_2681091 |
| MRPS18B | ProteinTech | Cat# 16139-1-AP<br>RRID: AB_2146368 |
| MRPS18C | Sigma | Cat# HPA050404<br>RRID: AB_2681115 |
| MRPS22 | ProteinTech | Cat# 10984-1-AP<br>RRID:AB_2146488 |
| MRPS25 | Sigma | Cat# HPA043490<br>RRID: AB_10794888 |
| MRPS25 | ProteinTech | Cat# 15277-1-AP<br>RRID: AB_2180358 |
| MRPS27 | ProteinTech | Cat# 17280-1-AP<br>RRID: AB_2180510 |
| NDUFA9 | Abcam | Cat# ab14713<br>RRID:AB_301431 |
| NDUFS1 | Thermo Fisher Scientific | Cat# PA5-22309<br>RRID: AB_11151879 |
| NFS1 | Proteintech | Cat# 15370-1-AP<br>RRID: AB_2151933 |
| NFU1 | Boster Bio | Cat# A06191<br>Validated in silenced cells |
| NFU1 | Abbexa | Cat# abx006894 |
| NUBPL | Abcam | Cat# ab171741 |

| | | |
|---|---|---|
| | | Validated in silenced cells |
| SDHA | Abcam | Cat# ab14715<br>RRID: AB_301433 |
| SDHB | Thermo Fisher Scientific | Cat# 459230<br>RRID: AB_2532233 |
| UQCRC2 | Abcam | Cat# ab14745<br>RRID: AB_2213640 |
| UQCRFS1 (Rieske) | Abcam | Cat# ab14746<br>RRID: AB_301445 |
| 2° Ab-mouse | Rockland Immunochemicals | Cat# 610-103-121<br>RRID: AB_218457 |
| 2° Ab-rabbit | Rockland Immunochemicals | Cat# 611-1302<br>RRID: AB_219720 |
| Chemicals, Peptides, and Recombinant Proteins | | |
| Dulbecco's Modified Eagle Medium (DMEM) | Invitrogen | Cat# 31966-047 |
| Fetal bovine serum (FBS) | Sigma | Cat# 12303C |
| Ham's F-10 Nutrient Mix, HEPES | Thermo Fisher Scientific | Cat# 12390035 |
| Lipofectamine RNAiMAX | Invitrogen | Cat# 13778150 |
| Endofectin | GeneCopoeia | Cat# EF014 |
| Opti-MEM™ I Reduced Serum Medium | Thermo Fisher Scientific | Cat# 31985062 |
| Digitonin, High Purity | Calbiochem | Cat# 300410 |
| Endothelial Cell Growth Kit-VEGF | ATCC | Cat# PCS-100-041 |
| $H_2O_2$ | Sigma | Cat# H1009 |
| Iodoacetamide | Sigma | Cat# I1149 |
| tris(2-carboxyethyl)phosphine | Invitrogen | Cat# T2556 |
| 4-Acetamido-4'-Maleimidylstilbene-2,2'-Disulfonic Acid, Disodium Salt | Invitrogen | Cat# A485 |
| Critical Commercial Assays | | |
| SuperSignal™ West Femto Maximum Sensitivity Substrate | Thermo Fisher Scientific | Cat# 34095 |
| Experimental Models: Cell Lines | | |
| HEK293T | ATCC | Cat# CRL-3216<br>RRID: CVCL_0063 |
| Primary Pulmonary Artery Endothelial Cells; Normal, Human (HPAEC) | ATCC | Cat# PCS-100-022 |
| MRPS25 Knockout – 293T | Synthego | Cat# 435091-1 |
| Oligonucleotides | | |
| *bS16m*_pCMV6_F: AGATCTGCCGCCGCGATGGTCCACCTCACTACTCTCC | This paper | N/A |
| *bS16m*_ pCMV6_R: CGCGGCCGGCCGTTTTTATGTTTCTGTAGCCTCTGTATCT | This paper | N/A |
| *bS16m*_C26A_F: CCTGGGTGGCgccACCAATCGGC | This paper | N/A |

| | | |
|---|---|---|
| *bS16m*_C26A_R: GCAAGGCGGATGGTTAAG | This paper | N/A |
| *bS16m*_C26S_F: CCTGGGTGGCagtACCAATCGGC | This paper | N/A |
| *bS16m*_C26S_R: GCAAGGCGGATGGTTAAGTG | This paper | N/A |
| *bS16m*_C26D_F: CCTGGGTGGCgatACCAATCGGC | This paper | N/A |
| *bS16m*_C26D_R: GCAAGGCGGATGGTTAAG | This paper | N/A |
| *mS25*_pCMV6_F: AGATCTGCCGCCGCGATGCCCAT GAAGGGCCGC | This paper | N/A |
| *mS25*_pCMV6_R: CGCGGCCGGCCGTTTTTAGTCCT GGGCATCGGC | This paper | N/A |
| Delta5_pCMV6_F: TGTCGTAATAACCCCGCC | This paper | N/A |
| Delta5_pCMV6_R: TGTACATATTATGATATAGATACAA C | This paper | N/A |
| *mS25*_C139A+C141A_F: cgccGAAGTGGAAGGGCAGGTG | This paper | N/A |
| *mS25*_C139A+C141A_R: atggcCTCCCGCAGGCAGTACTT | This paper | N/A |
| *mS25*_C139S+C141S_F: ctctGAAGTGGAAGGGCAGGTG | This paper | N/A |
| *mS25*_C139S+C141S_R: atagaCTCCCGCAGGCAGTACTT | This paper | N/A |
| *mS25*_C139D+C141D_F: cgatGAAGTGGAAGGGCAGGTG | This paper | N/A |
| *mS25*_C139D+C141D_R: atatcCTCCCGCAGGCAGTACTT | This paper | N/A |
| *bS16m*-genomic-exon2-F: AAGGGAATTGTGCCGTCTGT | This paper | N/A |
| *bS16m*-genomic-exon2-R: ACTGTCCTGGAGCTGACTTC | This paper | N/A |
| *mS25*-genomic-exon2-F: TTAGGTTAGCTTGCAGTACAGGTT | This paper | N/A |
| *mS25*-genomic-exon2-R: ACTTTGGAAAAATTACCCCCATGG | This paper | N/A |
| siNT | Ambion | Cat# oligo 4390844 |
| si*BOLA3*-1 | Ambion | Cat# oligo s52324 |
| si*BOLA3*-2 | Ambion | Cat# oligo s52322 |
| si*GLRX5* | Ambion | Cat# oligo s27700 |
| si*ISCU* | Ambion | Cat# oligo s23909 |
| si*ISCA1*-1 | Ambion | Cat# oligo s37761 |
| si*ISCA1*-2 | Ambion | Cat# oligo s37762 |
| si*NFU1*-1 | Ambion | Cat# oligo s26039 |
| si*NFU1*-2 | Ambion | Cat# oligo s26041 |
| si*NUBPL*-1 | Ambion | Cat# oligo s37094 |
| si*NUBPL*-2 | Ambion | Cat# oligo s37095 |
| si*NFS1* | Ambion | Cat# oligo s17266 |
| *ND2*_F: ATCTTAGCATACTCCTCAA | This paper | N/A |

| | | |
|---|---|---|
| *ND2*_R: GGTGTTAGTCATGTTAGC | This paper | N/A |
| *ACTIN*_F: TACCGTGAACTGGAACAA | This paper | N/A |
| *ACTIN*_R: TCGCAATAGGCAAAGATG | This paper | N/A |
| Recombinant DNA | | |
| *MRPS16* Human Gene Knockout Kit | OriGene | Cat# KN405837 |
| *mS25* in pReceiver-B02 | GeneCopoeia | Cat# EX-V0553-B02 |
| *bS16m* in pReceiver-B02 | GeneCopoeia | Cat# EX-V1236-B02 |
| pCMV6-A-Entry-Hygro | Origene | Cat# PS100024 |
| *bS16m*-WT in pCMV6-A-Entry-Hygro | This paper | N/A |
| *bS16m*$^{C26A}$ in pCMV6-A-Entry-Hygro | This paper | N/A |
| *bS16m*$^{C26S}$ in pCMV6-A-Entry-Hygro | This paper | N/A |
| *bS16m*$^{C26D}$ in pCMV6-A-Entry-Hygro | This paper | N/A |
| *mS25*-WT in Δ5-pCMV6-A-Entry-Hygro | This paper | N/A |
| *mS25*$^{C139AC141A}$ in Δ5-pCMV6-A-Entry-Hygro | This paper | N/A |
| *mS25*$^{C139SC141S}$ in Δ5-pCMV6-A-Entry-Hygro | This paper | N/A |
| *mS25*$^{C139DC141D}$ in Δ5-pCMV6-A-Entry-Hygro | This paper | N/A |
| *MRPS25-Flag_MRPS16-HA-His* in pETDuet-1 | GenScript | Cat# U0303FH310-4 |
| Software and Algorithms | | |
| ImageJ | NIH | https://imagej.nih.gov/ij/ |
| GraphPad Prism | GraphPad Software v.5.0a | N/A |
| Scaffold | Proteome Software, Inc | N/A |
| PyMol | Schrödinger, Inc. | N/A |
| Other | | |
